# Cell-subtype specific effects of genetic variation in the aging and Alzheimer cortex

**DOI:** 10.1101/2022.11.07.515446

**Authors:** Masashi Fujita, Zongmei Gao, Lu Zeng, Cristin McCabe, Charles C. White, Bernard Ng, Gilad Sahar Green, Orit Rozenblatt-Rosen, Devan Phillips, Liat Amir-Zilberstein, Hyo Lee, Richard V. Pearse, Atlas Khan, Badri N. Vardarajan, Krzysztof Kiryluk, Chun Jimmie Ye, Hans-Ulrich Klein, Gao Wang, Aviv Regev, Naomi Habib, Julie A. Schneider, Yanling Wang, Tracy Young-Pearse, Sara Mostafavi, David A. Bennett, Vilas Menon, Philip L. De Jager

## Abstract

The relationship between genetic variation and gene expression in individual brain cell types and subtypes has remained elusive. Here, we generated single-nucleus RNA sequencing data from the dorsolateral prefrontal cortex of 424 individuals of advanced age; analyzing 1.5 million nuclear transcriptomes, we assessed the effect of genetic variants on RNA expression in *cis* (*cis*-eQTL) for 7 cell types and 81 cell subtypes. This effort identified 10,004 eGenes at the cell type level and 8,138 eGenes at the cell subtype level. Many eGenes are only detected within cell subtypes. A new variant influences *APOE* expression only in microglia and is associated with greater cerebral amyloid angiopathy but not Alzheimer pathology, accounting for the effect of *APOEε4*, providing mechanistic insights into both pathologies. While eQTLs are readily detected, only a *TMEM106B* variant robustly affects the proportion of cell subtypes. Integration of these results with GWAS highlighted the targeted cell type and likely causal gene within susceptibility loci for Alzheimer’s, Parkinson’s, schizophrenia, and educational attainment.

## Introduction

After creating a critical new foundation for the study of neurodegenerative and neuropsychiatric diseases, gene discovery studies are reaching a point of diminishing returns with increasing sample sizes. We are left with a slowly growing list of validated susceptibility loci and a much larger set of loci that have not yet reached a threshold of genome-wide significance but are likely to contribute to disease susceptibility. Reference epigenomic atlases generated from small numbers of individuals or cell lines have been helpful in prioritizing variants within each locus and suggest which tissue or cell type may be relevant for the effect of a given variant (*1*–*3*).

These reference data are complemented by mapping quantitative traits which link haplotypes or individual variants with the abundance of a molecular feature; such “quantitative trait locus” (QTL) studies leverage the inter-individual heterogeneity of human populations to map the genetic architecture of molecular function. Until now, this was largely limited to tissue-level molecular profiles given the need for moderate to large-scale studies to achieve reasonable statistical power and the limitation of having to purify individual cells (*4*, *5*) and of imperfect estimates of cell types from bulk deconvolution efforts (*6*, *7*). However, cytometry, for example, enabled the execution of targeted protein measures on individual cells (*8*, *9*), and cell sorting approaches enabled the large-scale profiling of rare cell types (*10*, *11*), illustrating a path forward. Thus, to date, the mapping of quantitative traits remains largely limited to (i) bulk tissue profiles where traits such as RNA expression are averaged across all of the component cells of the tissue, (ii) limited dimensionality from the use of targeted molecular measures, or (iii) selected cell subtypes purified in isolation from the other cells in their tissue.

Single cell approaches have tantalized the human genetic community by their promise of providing a moderate throughput platform to produce single cell expression and epigenomic data. An early effort of modest size collating different small single nucleus-derived datasets from different brain regions and collections has shown that QTL mapping in this context is likely to be fruitful (*12*). Here, we generated a large set of data derived from individual nuclei extracted from the dorsolateral prefrontal cortex (DLPFC) of 424 older individuals sampled from a single, prospectively collected brain autopsy cohort. With these data in hand, we map *cis*-expression QTL in the 7 major cell types of the DLPFC, a neocortical region that is a hub for cognitive and mood circuits. Further, we map such effects within 81 cell subtypes, discovering additional *cis*-eQTLs influencing the expression of individual genes only in a subtype of cells, highlighting the importance of cellular context for uncovering many variants which influence gene expression.

This importance of nuanced gene expression is critical to understand, as efforts such as the Alzheimer’s Disease Functional Genomic Consortium get underway to map the functional consequences of disease-associated variation and study susceptibility genes in model systems. We thus go on to evaluate the transition of neocortical *cis*-eQTLs to induced pluripotent stem cell (iPSC)-derived neuronal (iN) and astrocytic (iAstro) model systems and find a set of 234 and 121 eGenes that replicate in iN and iAstro experimental systems, respectively. We also map the effects of genetic variants on the frequency of cell subtypes and evaluate the heritability of such fraction QTL (fQTL). Integrating our results with those of gene discovery studies, we prioritize (i) a set of cell types in which individual susceptibility loci appear to be having an effect despite wide expression of the target gene in a variety of cell types, (ii) variants in each risk haplotype that are the most likely to be the causal variant, and (iii) genes that may be the target of each variant.

## Results

### Description of the dataset

Figure 1A shows a schema of our study. Frozen neocortical samples for this project were accessed from the brains of participants in two longitudinal studies of cognitive aging with prospective autopsies: the Religious Order Study (ROS) and the Memory Aging Project (MAP) (*13*). All participants are without known dementia at baseline and agree to annual clinical evaluation and brain donation at the time of study entry; the two studies were designed and are run by the same group of investigators, with a large core of identical ante-and post-mortem phenotypic data collection. Thus, they are designed to be analyzed jointly (*14*) and are referred to as “ROSMAP”.

**Figure 1.**
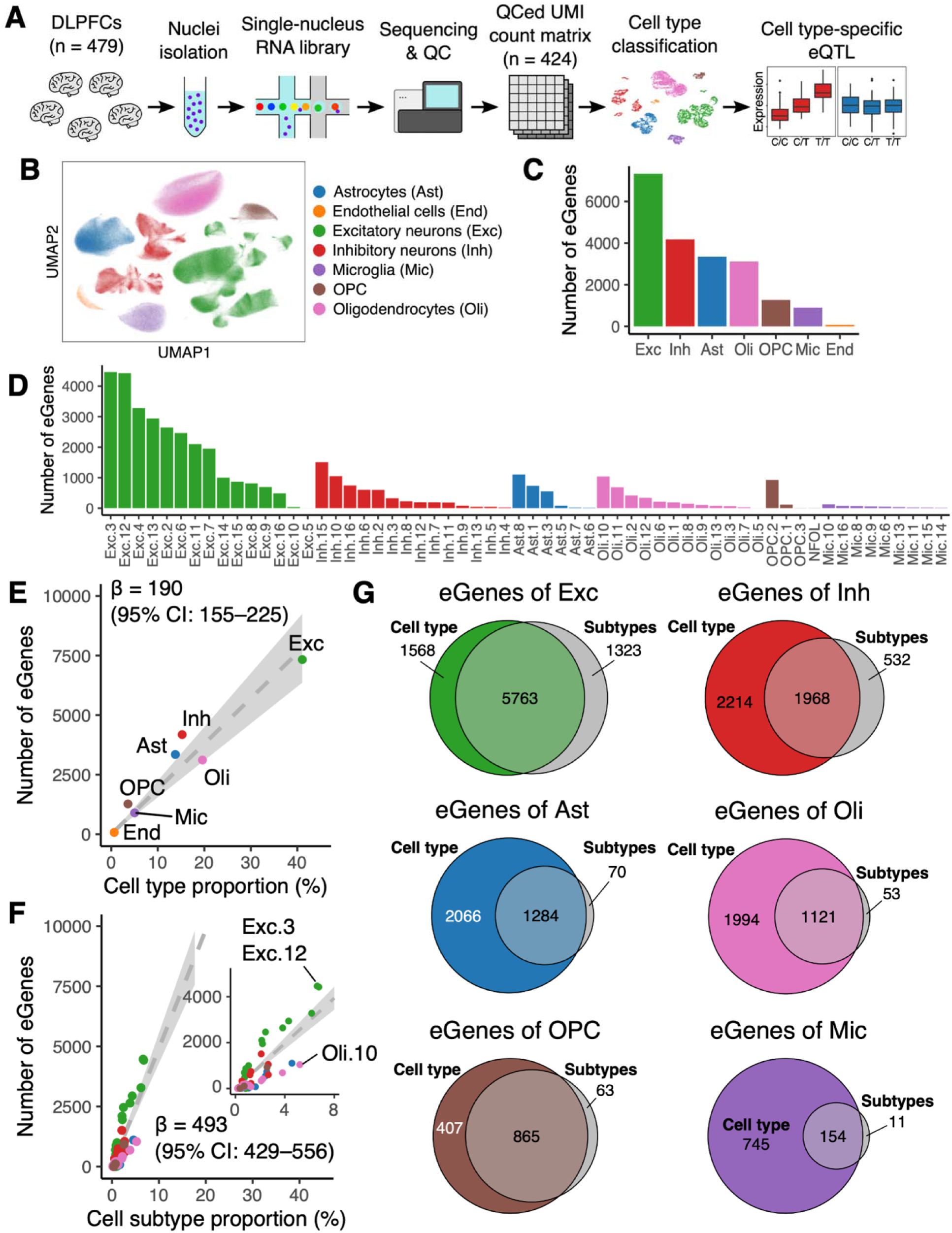
Study design and summary of cell type and subtype specific *cis*-eQTL. **(A)** Schema of our study. **(B)** UMAP visualization of 1,509,626 nuclei from 424 donors. Each of the seven major cell types is labeled with a different color. **(C)** Number of eGenes (genes targeted by a cis-eQTL effect) detected within each of the 7 cell types. **(D)** Number of eGenes detected in each of the 81 cell subtypes that were retained for analysis. **(E)** Relationship between cell type proportions and number of eGenes detected. β is a slope of linear fit. **(F)** Relationship between cell subtype proportions and number of eGenes detected. β is a slope of linear fit; the slope is much steeper than for cell types, as illustrated by the inset which enlarges the plotting of data near the origin. **(G)** Number of eGenes that are unique to the analysis of cell subtypes: for each cell type, we present a Venn diagram summarizing the extent to which cell type-level eGenes are found once the cells assigned to a given cell type are partitioned into the subtypes of that cell type; the six most common cell types are shown. For each cell type, the set of eGenes identified in all subtypes of a given cell type are shown in gray, and, in each cell type, a subset of these cell subtype eGenes are not recovered in the cell-type level analysis, suggesting that they may be specific to a cell subtype context.

Leveraging our prior work (*15*, *16*), single nucleus RNA sequencing (snucRNAseq) data was generated from a frozen fragment of the DLPFC; after thawing, the white matter was removed by dissection, and the gray matter was homogenized to create a nuclear suspension (**Methods**). Nuclei from 8 participants were pooled for data generation on the Chromium platform from 10X Genomics. The distribution of sex and clinical and pathological diagnosis among the 8 participants in each pool is shown in **Supplementary Figure 1**. All data completed a rigorous pre-processing pipeline, including assigning each nucleus to its source participant using RNA-derived genotypes and the Demuxlet method (*17*) (see **Methods** for details). In the frozen dataset, 424 participants with both snucRNAseq and whole genome sequence data were retained for analysis. Participants had a median of 3,824 sequenced nuclei. Demographic and diagnostic details of the 424 participants are presented in **Supplementary Table 1**. At the time of death, 34% of participants were cognitively non-impaired, 26% were mildly impaired, and 40% were demented. Of the 424 participants, 68% were female, and 63% fulfilled a pathologic diagnosis of Alzheimer’s disease (AD) by the NIH Reagan Criteria.

We used a stepwise clustering approach (**Methods**) to identify first the major cell types of the DLPFC and then subtypes in each cell type (Figure 1B and **Supplementary Figures 2 and 3**). In the end, we organized our data into 8 major cell types (7 of which had enough data for QTL mapping), which were further subdivided into 92 cell subtypes found in the human DLPFC.

**Figure 2.**
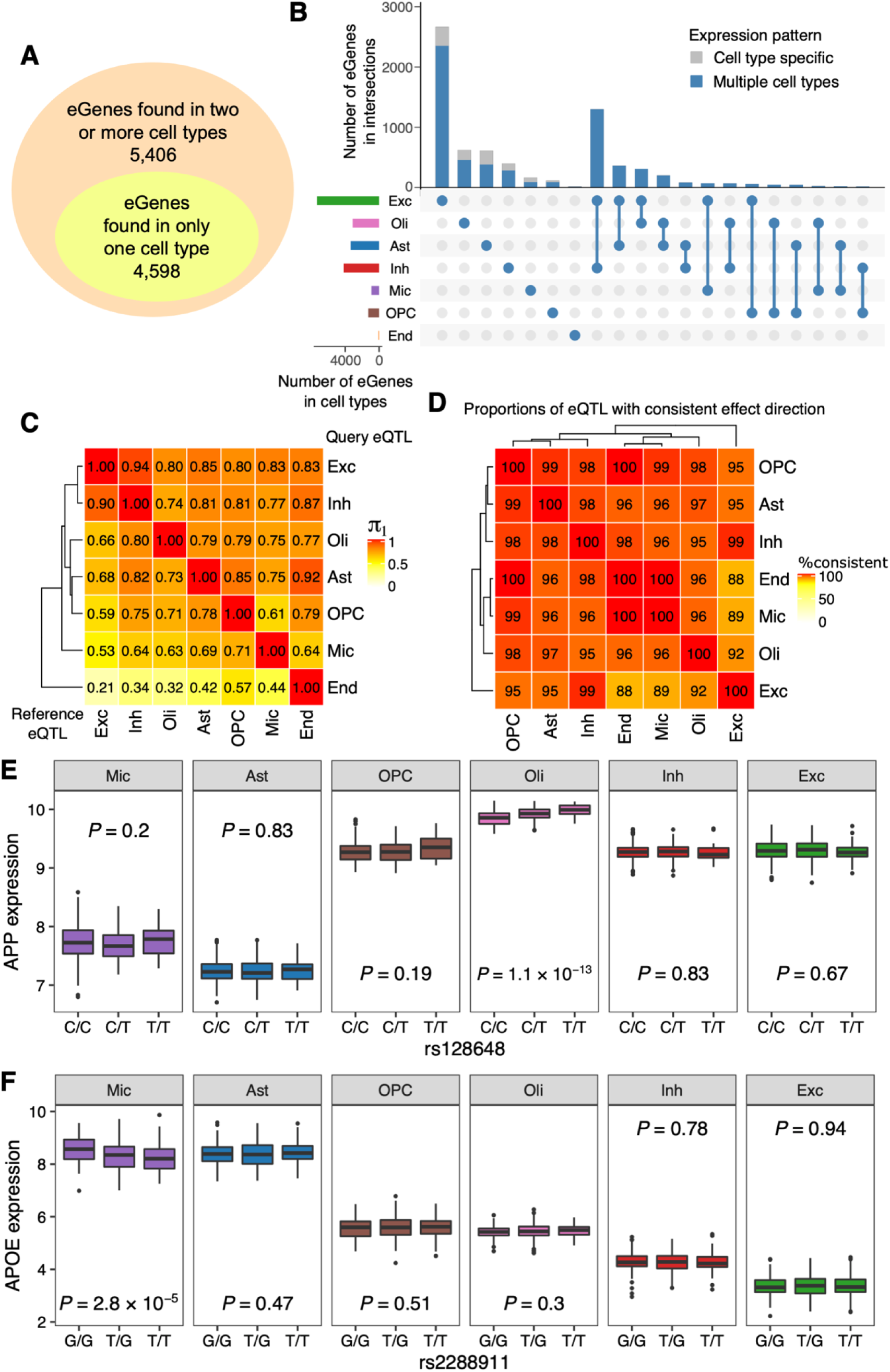
Similarities and differences of cell type-specific eQTL. **(A)** Number of cell type-specific and non-specific eGenes. **(B)** Number of eGenes that were unique to or shared between cell types. Only the top 20 intersections are shown. In the vertical bar chart, proportion of eGenes specifically expressed in one cell type are colored gray, and that expressed in two or more cell types are colored blue: most genes are expressed in more than one cell type. **(C)** π_1_ statistic to quantitate the extent of eQTL sharing between each pair of cell type. **(D)** Proportion of shared eQTL that had consistent direction of effect in each pair of cell types. **(E)** Example of an oligodendrocyte-specific eQTL found at the *APP* gene. *p*-values were computed by a simple linear regression between allele dosage of rs128648 and cell type-level expression of the *APP* gene. **(F)** Example of a microglia-specific eQTL at the *APOE* gene. *p*-values were computed by a simple linear regression between allele dosage of rs2288911 and cell type-level expression of *APOE* gene.

**Figure 3.**
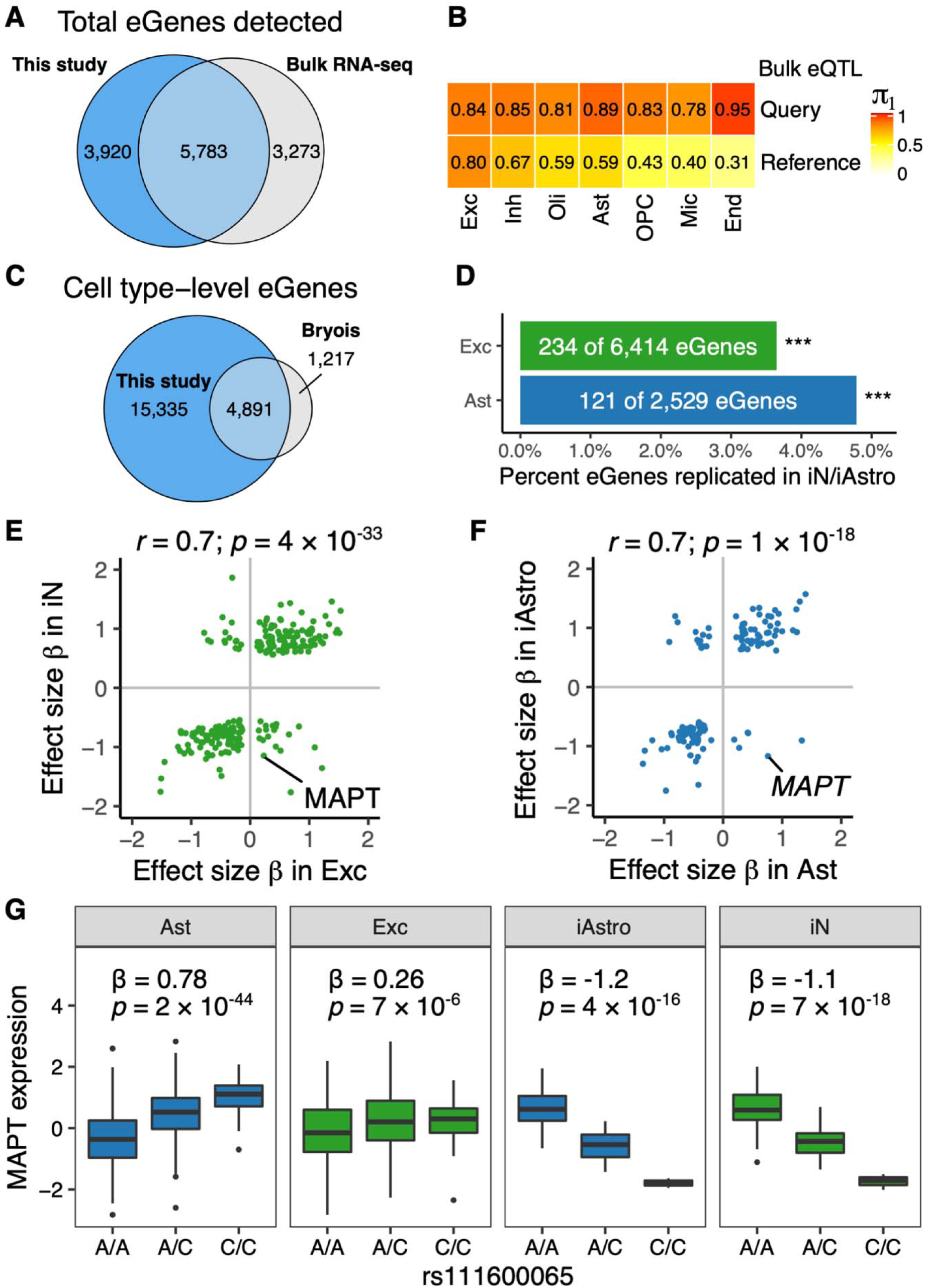
Replication of eQTL by bulk cortical RNA and RNA from induced cell lines. **(A)** Number of eGenes detected in our single-nucleus eQTL study vs. a bulk cortical RNA eQTL study using the same brain region (DLPFC). For the single-nucleus eQTL results, unique eGenes of the seven cell types were combined. For bulk cortical eQTL, we mapped *cis-*eQTL using 1,092 individuals from the ROSMAP studies. A large minority of eGenes are only found in the single nucleus data, despite the much smaller sample size. **(B)** π_1_ statistic of single-nucleus eQTL and bulk eQTL. The top and bottom rows used bulk eQTL as query and reference, respectively. As expected, lower frequency cell types are less likely to have their eGenes discovered in bulk cortical data. **(C)** Comparison of our single nucleus-derived eGenes and those from an earlier study (*12*) that had a smaller sample size and combined a mixture of brain tissues to produce results only at the cell type level. Our analysis recovered the vast majority of the earlier eGenes and identified 3-fold more eGenes, even without counting the cell-subtype level results. **(D)** Number of cell type-level eGenes that were replicated in iPSC-derived neurons (iN) and astrocytes (iA) generated from participants in the ROSMAP cohort. eGenes from excitatory neurons were tested in data from 44 iN lines, and astrocyte eGenes were tested in data from 38 iA lines, with each line coming from a different individual. In each cell type, the lead eSNP of each eGene was examined: we tested whether its allele dosage was associated with the eGene expression level in the corresponding iPSC-derived data with an FDR < 0.05. *** We replicate more of the eGenes than expected by chance, *P* < 1 × 10^-6^ by permutation test. **(E)** Effect size β of the 234 eGenes shared between excitatory neurons and iNs; most eGenes have a consistent direction and magnitude of effect, with some notable exceptions such as *MAPT*. **(F)** Effect size β of the 121 eQTL shared between astrocytes and iAstro. **(G)** Illustration of the opposite direction of effect for the *MAPT* eQTL: neurons and astrocytes display the same dosage effect at rs111600065, but the effect is reversed in the two iPSC-derived contexts. Effect size β and *p*-value were computed by linear regression between allele dosage of rs111600065 and *MAPT* expression. The major “A” allele tags the H1 haplotype of *MAPT*, and the minor “C” allele tags the H2 haplotype, associated with Parkinson’s disease, Parkinsonism, and perhaps AD (*63*, *64*).

### Mapping *cis*-eQTLs at the cell type level

To map *cis*-eQTLs, we created pseudo-bulk RNA expression measures for each major cell type by collapsing UMI counts from all nuclei assigned to a given cell type in each individual (see **Methods** for a detailed description of parameters). Separately, we repeated the process at the cell subtype level. To calculate eQTLs in a given cell type or subtype, we excluded donors with <10 nuclei assigned to the target cell type/subtype. We also excluded those cell subtypes with <10 participants with usable data. We used the Matrix eQTL version 2.3 method (*18*) method with 30 expression principal components, as we empirically determined this number as maximizing eQTL discovery (see **Methods**). After inclusion of these covariates, clinicopathologic traits and post-mortem interval did not meaningfully influence eQTL discovery. **Supplementary Figure 4** shows the number of genes tested for eQTL in each cell type. **Supplementary Table 2** summarizes the results of *cis*-eQTL mapping at the cell type level. In Figure 1C, we summarize the number of eGenes detected, where an “eGene” is a gene for which a SNP:gene pair exceeds our threshold of significance, a two-step FDR < 0.05 (**Methods**). Using a similar approach, Figure 1D and **Supplementary Table 3** present the number of eGenes detected in the 81 of our 92 cell subtypes retained for eQTL analysis.

The large difference in the number of eQTLs between cell types is explained in part by cell type proportions (Figure 1E). Given that neurons – particularly the excitatory type but also true for inhibitory neurons – are the most abundant cell types in the neocortex and also have, on average, the most RNA, it is not surprising that they drive the largest proportion of eGene discovery. On the other hand, less common cells such as microglia – which are, on average, 5.0% of nuclei in a participant and are known to have a relatively small amount of RNA – return a substantial (899 eGenes) but smaller number of eGenes when compared to excitatory neurons (7,331 eGenes). The correlation between cell population frequency and eGene discovery also holds among cell subtypes (Figure 1F). Unexpectedly, the slope of eGenes per cell population proportions was much steeper among cell subtypes than in cell types (respectively β = 493 and 190 per 1% in cell subtype/type frequency; *p* = 1.3 × 10^-12^), suggesting that cell subtypes may be a better target for eGene discovery in future studies.

To evaluate the extent to which cell subtype analysis enhances *cis*-eQTL discovery, we compared eGenes between each cell type and its subtypes (Figure 1G). In excitatory neurons, 1,323 unique eGenes were only detected in subtypes of excitatory neurons but not in the top-level cell type in which all excitatory neuron subtypes are pooled. The biggest gain comes from the neuronal subtypes in part because neurons are abundant and rich in RNA content, but perhaps also because they contain more discrete cell subtypes. Nonetheless, there is a non-negligible gain of eGenes when analyzing subtypes for all other subtypes, including astrocytes, microglia, oligodendroglia, and oligodendroglial progenitor cells (OPCs) (where the boundaries defining cell subtypes are less stark than among neurons) (Figure 1G).

### Similarity and specificity of neocortical eGenes

Among the 10,004 eGenes that we have detected across all cell types, the majority (5,406 eGenes; 54%) were present in more than one cell type, but we find that a substantial proportion (4,598 eGenes; 46%) is cell-type specific as well (Figure 2A). In Figure 2B, we illustrate the specificity of *cis*-eQTLs by displaying the extent to which they are shared among the different cell types. For example, there are 622 oligodendrocyte-specific eGenes, and, for microglia that are implicated in several CNS diseases, we found 165 eGenes which are specific to this cell type.

Cell-type specificity of eQTL can be explained either by (i) a target gene that is expressed only in one cell type, or (ii) a target gene that is expressed in multiple cell types but that has the genetic association only in one cell type. Interestingly, a large fraction of cell type-specific eGenes fit the latter pattern (blue bars in Figure 2B) rather than the former (gray bars): the majority of cell type-specific eGenes may thus be driven by context-specific regulators of gene expression and not master regulators of cell type differentiation. A similar trend was observed in cell subtype analysis (**Supplementary Figure 5)**.

To assess the extent of eQTL sharing between cell types, we computed a π_1_ statistic (Figure 2C). Aside from endothelial cells, which had a limited number of eGenes given their low frequency among the nuclei, the six other cell types had a high degree of eGene sharing (π_1_, 0.53–0.94). For those *cis*-eQTLs which are shared, Figure 2D shows that the vast majority of shared eQTLs have the same direction of effect, and **Supplementary Figure 6** shows that the effect size β is also strongly correlated between any pairs of cell types. However, there are a number of cell type specific effects. Figure 2E shows one example at *APP*, an important gene related to Alzheimer’s disease (AD): *APP* expression is associated with rs128648 only in oligodendrocytes. Figure 2F presents another instance of cell-type specificity as *APOE* is expressed in 6 different cell types but only shows a novel *cis*-eQTL effect in microglia, where eQTL detection is relatively underpowered compared to other cell types. This is interesting as *APOE* is one of the key genes upregulated in Disease Associated Microglia (DAM) that have been described in amyloid proteinopathy models (*19*). This *APOE* locus variant, rs2288911, is associated with AD according to summary statistics of previous GWAS (*p* = 1.0 × 10^-12^ and *p* = 6.0 × 10^-10^) (*20*, *21*). In analyses of AD-related neuropathological measures in ROSMAP cohort, this variant is not associated with amyloid or tau proteinopathy (*p* > 0.05). However, this variant is associated with the burden of cerebral amyloid angiopathy (CAA) (*p* = 1.18 × 10^-7^) (**Supplementary Figure 7**), and this association persists even after accounting for the effect of the *APOE* ε4 haplotype on CAA (*p* = 9.9 × 10^-6^). There is no evidence of interaction between the effects of rs2288911 and *APOE* ε4 on CAA burden (*p* > 0.05). Thus, this eQTL appears to be independent of the well-validated *APOE* ε4 risk haplotype, which alters the coding sequence of APOE and does not affect *APOE* expression level. Thus, increased expression of APOE in microglia leads to more CAA but not AD pathology, causing microhemorrhages that contribute to dementia and probably explain the association to AD dementia. This may be similar to the association of the *GRN* locus with AD as *GRN* causes frontotemporal dementia and contributes to the mixed dementia of older age, along with neurovascular factors such as CAA.

### Comparison to existing datasets of bulk cortex, purified cell types, and an earlier single nucleus effort

One way to assess the robustness of our results is to compare our SNP:Gene eQTL pairs to those that have been robustly established in related efforts. An important comparison in this regard involves the results of analyses using the same methods and thresholds in bulk RNA-seq data from the DLPFC of ROSMAP participants: our earlier *cis*-eQTL mapping in this context (and subsequent efforts) reported good agreement of our *cis*-eQTLs with efforts in other sample collections using the same or similar bulk neocortical tissue (*5*, *22*, *23*). Here, we expanded our earlier effort (*5*) and mapped *cis*-eQTLs from bulk RNA-seq of the same DLPFC region from 1,092 ROSMAP participants. 408 participants are shared between the bulk and snucRNAseq analyses. **Supplementary Table 4** presents a summary of the bulk *cis*-eQTL mapping. In Figure 3A, we illustrate that the majority of eGenes are detected in both datasets; the bulk-specific effects are not surprising given the greater sample size and greater complexity of the bulk transcriptome (which contains cytoplasmic RNA) relative to the sparse transcriptomes of individual nuclei. However, it is striking that 40% of snucRNAseq-derived eGenes are not discovered in the larger bulk cortical RNA-seq dataset from the same brain region, confirming the importance of mapping eGenes within cell types and subtypes. Figure 3B and **Supplementary Figure 8** present the results of the π_1_ statistics for each cell type; as expected, sharing is greatest in those cell types that are more abundant and express more RNA in the neocortical parenchyma, resulting in a greater contribution to the bulk RNA-seq transcriptome. Microglia, on average 5.0% of the nuclei in this region, contribute only a small fraction of the DLPFC transcriptome, and thus its more specific expression patterns are swamped in the homogenization of the RNA that occurs in the bulk RNA preparation.

Another important comparison in detecting eGenes involves contrasting our snucRNAseq-based results with those from a purified cell type profiled by bulk RNA-seq since the latter effort will capture the cytoplasmic transcriptome. We therefore compared our microglial results from snucRNAseq to a recently published set of results from bulk microglia profiles generated from human microglia isolated from autopsy tissue (*11*). **Supplementary Figure 9** shows the modest overlap between the two datasets of microglial eGenes. The limited overlap may be partly explained by the single-nucleus nature of this study, which cannot detect cytoplasmic components and the small sample size of the bulk microglial profiles (n = 77 medial frontal gyrus).

Finally, we also compared our results to those recently generated from the merger of several small snucRNAseq datasets from different brain regions (*12*). In these analyses, we validate many of the results of the earlier, smaller effort: 80% of cell type:eGene pairs in the previous study are included in our results (Figure 3C). The π_1_ statistics is 0.90 when the results of the prior manuscript were used as reference. Further, we also illustrate the impact of targeting a single neocortical region and a larger sample size, as we have now uncovered 15,335 new eGene: cell type pairs, and, importantly, we have now mapped eGenes at the cell subtype level, which the earlier effort did not.

### Translating results *in vitro* to guide functional studies of genetic variants

With the vast number of disease susceptibility variants identified to date, an urgent challenge involves mapping their functional consequences in human tissues but also in model systems such as iPSC-derived cells, where manipulation of the genome or targeted perturbations of the cellular culture environment can occur. As noted above, most of our eGenes are expressed in multiple cell types but display the effect of the variant in only one or a subset of the neocortical cell types; thus, simply screening for gene expression is inadequate when selecting a cell type of interest for an iPSC model. To address this issue, we selected two common model systems – iPSC-derived neurons (iN) and astrocytes (iA) – and evaluated whether they displayed the same eGenes identified from our snucRNAseq data. Specifically, we repurposed separate bulk RNA-seq from iN (n= 44) and iAstro (n=38) cultures; each of these iPSC lines was derived from a different participant in the ROSMAP cohort and have whole genome sequence data. The iN have certain amyloid-related *in vitro* phenotypes that are associated with brain-derived traits, making them interesting proxies for mechanistic dissection (*24*). Given the modest sample size for iPSC-derived eQTL analysis, we limited our evaluation to replicating the excitatory neuron and astrocytic eGene results rather than *de novo* discovery of eGene. Results are shown in Figure 3D: several hundred eGenes identified in the neocortical nuclei are replicated in the iN’s and in iAstro, with a slightly larger proportion of iAstro eGenes replicating, perhaps because astrocytes show less inter-subtype diversity. Specifically, 6,414 genes were expressed in iNs as well as in our excitatory neuron cell type pseudobulk data, and expression levels of 234 genes (3.6% of assessed eGenes) were associated with eSNPs at an FDR < 0.05. A permutation test confirmed that the observed associations are significantly more frequent than random pairs of genes and SNPs (*P* < 1 × 10^-6^). For astrocytes, 121 of 2,529 (4.8%) eGenes were reproduced in iAstro (*P* < 1 × 10^-6^). Larger efforts generating iN, iAstro, and other cell types such as iMicroglia will be needed to comprehensively evaluate the translatability of our eGenes; however, this initial analysis already shows that many effects will be reproduced in an artificial, *in vitro* context. The effect sizes are largely similar, and most are in the same direction (snuc vs. iN/iAstro data) (Figure 3E and 3F). However, there are some interesting exceptions, including the *MAPT* gene, which encodes the tau protein. As seen in Figure 3G, the rs11100065 variant is significant in all 4 contexts, but the direction of the effect on *MAPT* expression is inverted in the iPSC-derived contexts. Given the strength of the effect *in vitro*, its presence in both iN and iAstro and the fact that most eGenes have an effect in the same direction (snuc vs. iPSC-derived), this is unlikely to be a statistical fluctuation. Further work will be needed to resolve the nature of this effect, particularly as two other studies of bulk neocortical data have reported a direction of effect similar to the iPSC-derived cells (*25*, *26*). Nonetheless, it is already a cautionary tale that even well-studied but genetically complex loci like the one harboring *MAPT* can harbor substantial surprises when being studied in a reductionist system that by its nature eliminates many additional factors such as age that may interact with genetic variants to affect expression levels.

### Chromatin annotation of *cis*-eQTLs

To further characterize and validate our *cis*-eQTL results, we annotated them with available epigenomic features from bulk DLPFC (*1*) and relevant cell types; the latter are generated as bulk profiles from 4 different cell types isolated from cerebral cortex (*27*). As seen in Figure 4A, we see the expected enrichment of our *cis*-eQTL SNPs within ∼100 kb of the target gene’s transcription start site (TSS), which is the typical pattern for *cis*-eQTLs (*28*, *29*). Further, these SNPs are enriched in DLPFC euchromatin (transcriptionally active regions and enhancers) and relatively depleted in heterochromatic regions (Figure 4B). Looking more closely at euchromatin captured from bulk profiles from four purified cell types (Figure 4C), we see larger enrichment among TSS. Still, we also see a clear enrichment among those enhancers annotated in the corresponding cell type. For example, SNPs for microglial eGenes are more likely to be found in enhancer regions found in bulk microglia than in neuronal, oligodendroglial, or astrocytic enhancers. In Figure 4D, we show the distribution of peaks from chromatin immunoprecipitation followed by sequencing (ChIP-seq) in the *APOE* locus relative to the rs2288911 SNP that drives a microglial-specific *cis*-eQTL of *APOE* (Figure 2F). This SNP is 40.2 kb from *APOE* (near another gene, *APOC2*) but in a chromosomal segment decorated, only in microglia, with H3K27ac and H3K4me3, two marks associated with active transcription and enhancers. Further, PLACseq data suggest that this chromosomal segment is in physical proximity to the *APOE* gene, which is also in an active conformation. Thus, it is plausible that this variant may be driving the observed effect on microglial *APOE* expression, while the astrocytic *APOE* locus (which is also in a transcriptionally active conformation) is unaffected by rs2288911, suggesting that microglia may play a key role in the accumulation of CAA described above by the risk allele at this SNP.

**Figure 4.**
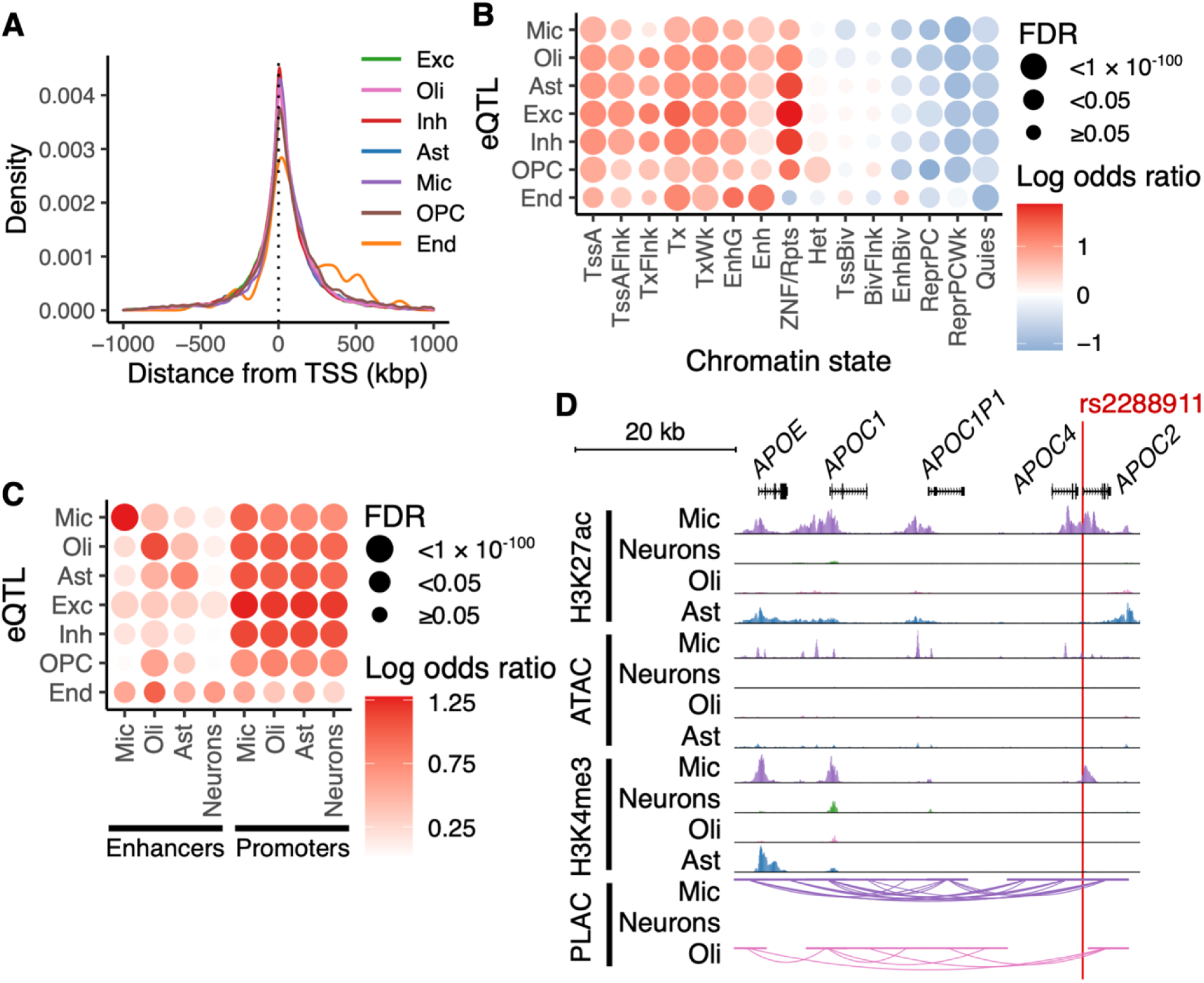
Chromatin states of cell type-level eQTL. **(A)** Distance between eSNPs and transcription start sites (TSSs) of eGenes. **(B)** Enrichment of eSNP in chromatin states. Chromatin states of a human DLPFC tissue were obtained from sample E073 of Roadmap Epigenomics (*1*). TssA, active TSS; TssAFlnk, flanking active TSS; TxFlnk, transcription at gene 5’ and 3’; Tx, strong transcription; TxWk, weak transcription; EnhG, genic enhancers; Enh, enhancers; ZNF/Rpts, ZNF genes & repeats; Het, heterochromatin; TssBiv, bivalent/poised TSS; BivFlnk, flanking bivalent TSS/enhancer; EnhBiv, bivalent enhancer; ReprPC, repressed PolyComb; ReprPCWk, weak repressed PolyComb; Quies, quiescent/low. **(C)** Enrichment of eQTL in cell type-specific enhancers and promoters. Enhancers and promoters of four brain cell types were obtained from a published report (*27*) **(D)** Microglia-specific eQTL for *APOE* (rs2288911) in the context of microglial-specific chromatin conformation data. These data were repurposed from Nott et al. (*27*). The x-axis denotes the physical position along a segment of chromosome 19 containing the APOE gene and several related genes; their exon structure is presented in the top horizontal track. The next four tracks report chromatin immunoprecipitation followed by sequencing (ChIP-seq) data against the H3K27Ac epitope, a mark found in active TSS and enhancers; each track presents data from a different cell type, isolated as purified nuclei. The peaks denote segments that are in a transcriptionally active conformation. The next four tracks present data from the same samples using the Assay for Transposase Accessible Chromatin (ATAC) which denotes chromosomal segments that in an open conformation and accessible for transcription. The four H3K4Me3 tracks present ChIP-seq data for that epitope, which is also correlated with transcriptionally active regions of a chromosome. Proximity ligation-assisted ChIP-seq (PLAC-seq) data are presented in the last 4 tracks and denote pairs of chromosomal segments that are in physical proximity to one another as the chromatin loops.

### Identifying variants influencing the proportions of cell subtypes

Having evaluated the effect of genetic variation on the expression of individual genes in *cis*, we also performed genome-wide studies to identify variants influencing the proportion of each cell subtype: this is an effort to discover fraction quantitative trait loci (fQTL). Results are presented in Figure 5A and **Supplementary Table 5**. We find 44 loci that meet our significance threshold, *p* < 5.6 × 10^-10^. This threshold was determined by applying Bonferroni correction for 90 tested cell subtypes to the standard genome-wide threshold of p < 5 × 10^-8^. The distribution of associations is not uniform, suggesting that some subtypes may be under stronger genetic influence than others. To assess this possibility, we calculated the SNP-based heritability of each of the frequencies of cellular subtypes that we evaluated. This analysis returns only four subtypes with modest evidence of heritability: excitatory neuron 10, oligodendrocyte 10, endothelial cell 1, and fibroblast 1 (**Supplementary Table 6**). The statistical power of our dataset may not be sufficient to evaluate the heritability of cell subtype proportions, but these results suggest that the frequency of many of these subtypes may not be strongly influenced by genetic variation. Nonetheless, selected results may well be robust, particularly those in which the fQTL SNP is also a *cis*-eQTL, as in the case of the loci shown in **Supplementary Table 7**.

**Figure 5.**
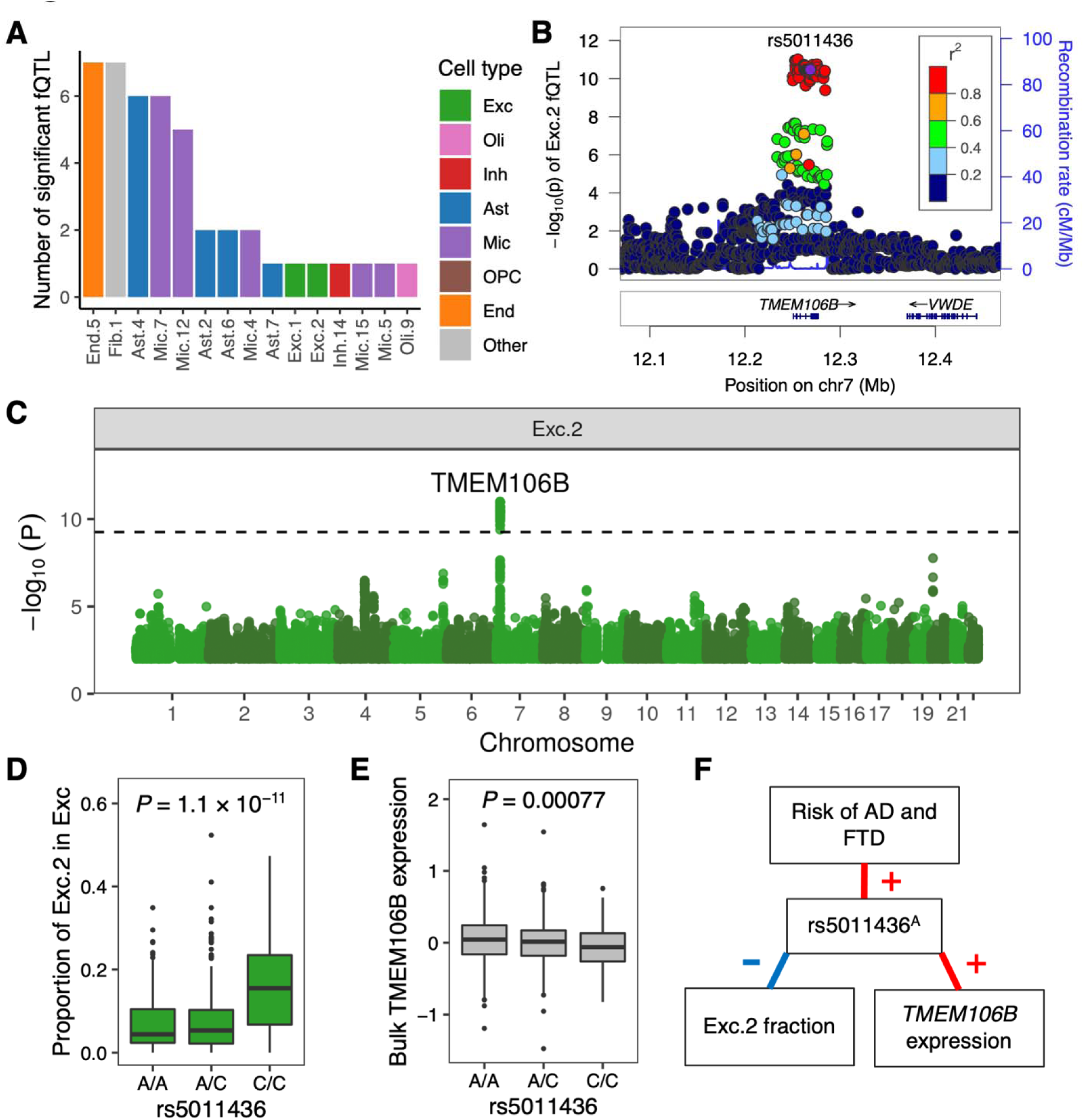
Fraction QTL (fQTL) between SNPs and cell subtype proportions. **(A)** Number of independent significant fQTL per cell subtypes. **(B)** Locuszoom plot around the *TMEM106B* locus. A lead risk SNP of AD in the locus (rs5011436) was used as the reference SNP given that it tags a risk haplotype that contains many other SNPs with similar statistical properties given their strong LD (*30*). The *y*-axis shows-log_10_ of *p*-values for the Excitatory neuron subtype 2 (Exc.2) cell subtype fQTL. **(C)** Manhattan plot for genome-wide fQTL results for Exc.2. The dashed line shows the significance threshold *p* < 5.6 × 10^-10^ (accounting for al tested fQTL GWAS). **(D)** Genotypes of rs5011436 and proportion of Exc.2 among excitatory neurons. **(E)** Genotypes of rs5011436 and gene expression levels of *TMEM106B* in bulk DLPFC tissues, illustrating the local cis-eQTL. **(F)** Schematic summary of the major allele of rs5011436 and its relationship to three phenotypes. FTD, frontotemporal dementia. Red and blue lines show positive and negative correlations, respectively.

We then examined relevance of fQTL to Alzheimer’s disease (AD) risk. Among AD risk SNPs we tested (*30*), only one SNP (rs5011436) coincided with fQTL of excitatory neuron 2 (Exc.2) (Figure 5B). This SNP in the *TMEM106B* locus and was the sole fQTL for Exc.2 (Figure 5C). The SNP not only is an fQTL but also has a cis-eQTL effect on the expression of *TMEM106B* in bulk cortical RNAseq data (Figure 5D–F). The proportion of inhibitory neuron 6 (Inh.6) also had a marginal association with this SNP, although it was below the genome-wide significance threshold (**Supplementary Figure 10**). The major allele rs5011436^A^ is a risk allele for AD (*21*, *30*), but this association may relate to its role as a tag for the susceptibility haplotype for frontotemporal dementia (*31*), which may be why this locus emerges in certain AD GWAS. In addition, we ran a meta-PheWAS for this variant using the United Kingdom Biobank data and the eMERGE-III data (*32*, *33*); interestingly, the two significant results out of 1,817 clinical traits tested were related to diabetes (**Supplementary Figure 11, Supplementary Table 8**), and three of the top five suggestive results (*p* < 0.001) relate to atherosclerosis and include cerebrovascular disease. This is notable as both diabetes and vascular disease influence risk of AD and dementia: this TMEM106B variant may therefore influence certain neuronal proportions through vascular/metabolic effects.

Using the CelMod method (*16*), we inferred the proportion of all cell subtypes using bulk DLPFC RNASeq data from 1,092 ROSMAP participants and retained those subtypes imputed with high confidence (see **Methods**). Using these imputed cell type frequencies, we replicated one of the 44 fQTL effects discovered in the snucRNAseq data. It was the effect of rs150928213 on the Ast.2 subtype fraction (nominal *p* = 2.1 × 10^-5^). The SNP is in an intronic region of the Neuron Navigator 1 (*NAV1*) gene but was not a *cis-*eQTL in astrocytes. In this CelMod analysis, the Exc.2 fQTL with rs5011436 did not replicate (nominal *p* = 0.39) but that of the Inh.6 neurons with rs5011436 became significant (nominal *p* = 5.0 × 10^-9^).

### Colocalization with disease susceptibility

An important use of our *cis*-eQTL results involves aligning them with the results of gene discovery studies to assess whether altered gene expression may be the mechanism for a particular risk allele or haplotype. We used the updated COLOC method (version 5.1.0) R package to conduct colocalization analysis for this purpose. Using a genome-wide association study of Alzheimer’s disease and dementia (AD) (*30*) and our list of eGenes, we find evidence of colocalization (PP.H4 > 0.8) for 21 eGenes among the 20 AD loci that we interrogated (Figure 6A). We confirm some of the well-validated results, such as the *BIN1* risk haplotype tagged by *rs4663105*, the susceptibility haplotype with the largest effect size for AD after *APOE,* which drives *BIN1* expression only in microglia (*11*); other cell types express BIN1 but don’t exhibit this effect. More interesting is that we find 4 novel colocalizations with our data: *AC004797.1*, *AL596218.1*, *AP001439.1*, and *ITGA2B*. As expected with AD, we find that microglia harbor the most implicated target genes, although all other neocortical cell types are also implicated by our colocalization analysis (Figure 6A). The endothelial cells have not been sequenced deeply (few nuclei per individual) in our participants and have very few eGenes so far. Therefore, we cannot interpret the lack of colocalization so far. We also note that while many loci have unambiguous cell type specific effects (i.e., *CASS4* or *ACE*), one has an effect shared by three cell types (*APH1B*: Ast, Exc. Neurons, and Oligo), and another has distinct target genes in two cell types (*CR1/AL5962.81* in Oligodendrocytes and OPCs). While strong posterior probabilities are seen in most loci, some loci have poor evidence so far (*ZCWPW1*) or muddled evidence of colocalization that will require further dissection (*SCIMP1*). As reported previously, we find colocalization in the *GRN* locus, but, interestingly, it appears to be found in both oligodendroglial cells and excitatory neurons. Further, there is some evidence that another gene, *ITGA2B*, may also be implicated (although modestly) in excitatory neurons.

**Figure 6.**
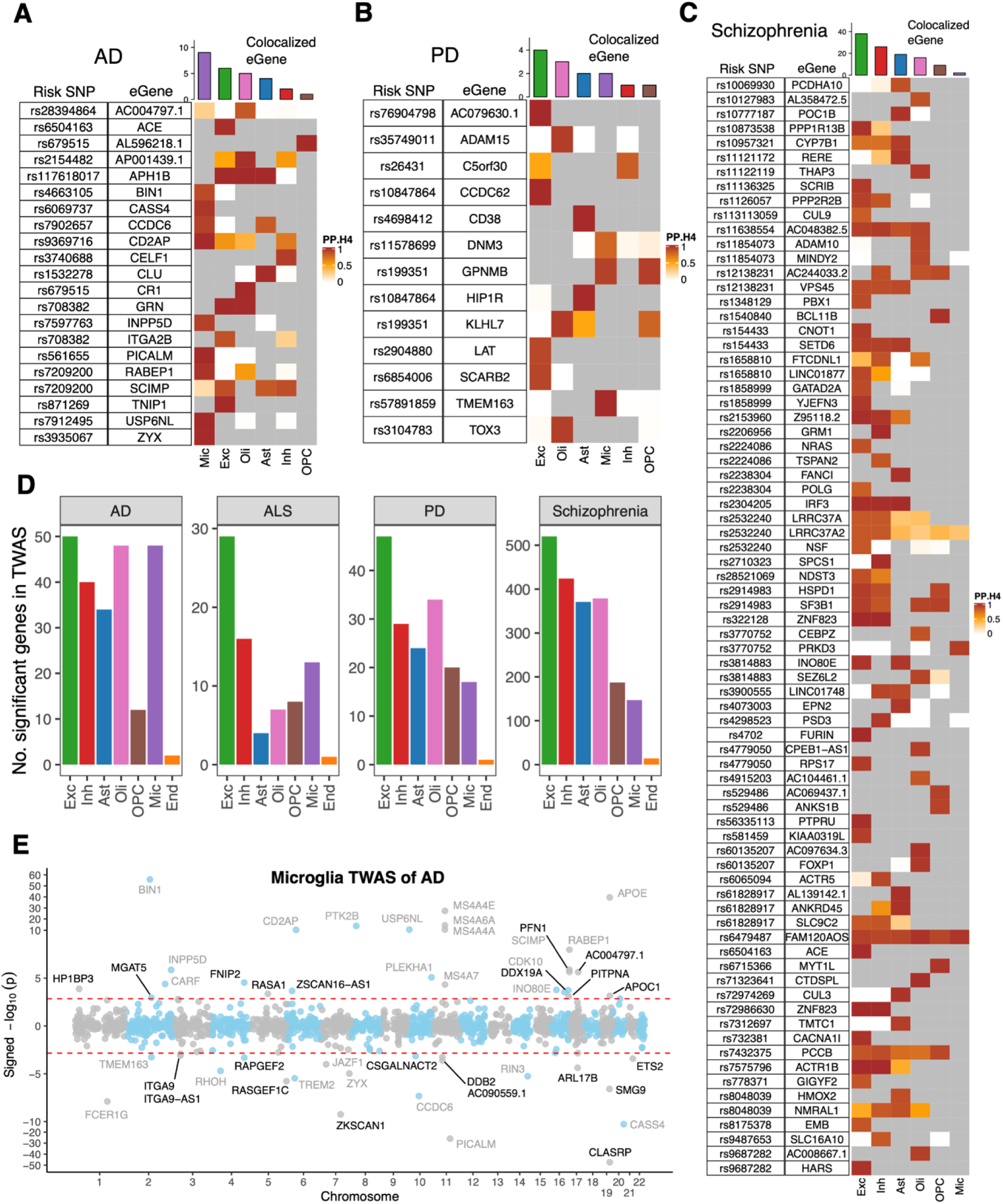
Overlap of results for our eQTL and GWAS of selected neurodegenerative and neuropsychiatric disease. **(A–C)** Colocalization of cell type-specific eQTL with risk SNPs of **(A)** Alzheimer’s disease GWAS (*30*), **(B)** Parkinson’s disease GWAS (*52*), and **(C)** Schizophrenia GWAS (*53*). Each of the heatmaps report the posterior probabilities of the H4 hypothesis (PP.H4) of the coloc method (*65*), which assumes GWAS and eQTL share a single causal SNP. Rows report overlap for individual genes and SNP pair; columns report PP.H4 score in each of our cell types. The color of each cell is based on the code found to the right of each panel; the darker color denotes higher confidence that the same variant influences susceptibility and gene expression in that cell type. Grey cells indicate that the gene was not an eQTL target in that cell type. Top bar chart shows the number of colocalized eGenes with high confidence (PP.H4 > 0.8) in each cell type. **(D)** Cell type-level transcriptome-wide association studies (TWAS). Using the FUSION method, we deployed instruments inferring the expression of 22,159 genes across all cell types in the summary statistics for AD, PD, ALS, and schizophrenia. The count of genes meeting a transcriptome-wide threshold of significance in each cell type is presented for each disease, with the expected excess of microglial genes in AD and intriguing number of oligodendroglial genes in PD. Cell types are in order of descending expression heritability. **(E)** Illustration of the TWAS result from one cell type in one disease: the statistical significance and effect direction of all inferred microglial genes are presented with the physical position along the chromosome being presented on the x-axis and the significance threshold, on the y-axis. Each dot represents a gene. Positive and negative *y* coordinates show that transcript abundance was associated with increased and decreased risk of AD, respectively. The *y*-axis between-10 and 10 are enlarged to enhance visibility. Novel and known candidates for AD risk genes in microglia are colored black and gray, respectively. Red dashed lines highlights the threshold of FDR = 0.05.

While the frontal cortex from aging and AD subjects may be most relevant to interpreting the results of AD GWAS, we illustrate the utility of our resource by conducting similar analyses for schizophrenia, a neuropsychiatric disease in which the frontal cortex is also implicated, and other neurodegenerative diseases such as amyotrophic lateral sclerosis (ALS) and Parkinson’s disease (PD). For the PD COLOC, we identified 11 loci, mapping to 13 eGene (Figure 6B). For the schizophrenia COLOC, we identified 57 loci and 75 eGenes (Figure 6C). For ALS, COLOC identified one locus and one eGene (**Supplementary Figure 12A**). For both PD and schizophrenia, we found that excitatory neurons achieved the largest number of colocalized loci. ALS and PD are diseases that strongly affect other CNS regions, but we can already prioritize several genes as being colocalized in specific cell types, which are shared among these CNS regions. The results of our colocalization analyses are summarized in **Supplementary Table 9.** In addition, we conducted COLOC analyses for educational attainment and brain volume (**Supplementary Table 10**): for educational attainment, we identified 16 colocalized loci and 21 eGenes, while, for intracranial brain volume, we identified 3 loci and 7 eGenes. Other MRI-derived volumetric traits include (1) cortical surface area and thickness with 3 loci and 2 eGenes; (2) accumbens volume with 1 locus and 3 eGene; (3) Brainstem had 5 loci and 5 eGene; (4) Caudate had 4 loci and 4 eGene; (5) Pallidum had 6 loci and 5 eGenes, (6) Putamen volume with 6 loci and 5 eGenes, (7) Thalamic volume with 2 loci and 2 eGenes. Finally, we found no colocalized loci for hippocampal volume.

### Prioritizing additional loci and genes as implicated in AD and other neuropsychiatric diseases

In our data, we find that the expression level of thousands of genes is strongly heritable in all cell types except endothelial cells which are under-sampled in our dataset (**Supplementary Figure 12B and Supplementary Table 11**). Here, heritability was assessed using the genomic-relatedness-based restricted maximum-likelihood (GREML) approach with a threshold of *p*-value ≤ 0.05 (*34*). This enables us to perform a “transcriptome-wide association study” (TWAS) for each cell type in which we infer gene-level association statistics by combining GWAS results for a trait and a gene’s model to infer RNA expression. This is complementary to the colocalization studies where individual variants and haplotypes are assessed; the TWAS is particularly helpful in prioritizing genes for further evaluation in loci where no individual variant has reached a threshold of genome-wide significance. In Figure 6D, we display summaries of TWAS for different diseases, confirming the large excess of microglial genes involved in AD relative to other neuropsychiatric diseases. In Figure 6E and **Supplementary Figure 13**, we illustrate the large number of positive results in the AD TWAS using microglial eGenes, highlighting *FNIP2* and *RAPGEF*2, two of 22 microglial genes which have not previously been associated with AD in similar analyses using other forms of RNA data. **Supplementary Table 12** shows details of TWAS results for neuropsychiatric diseases.

The TWAS analysis is also helpful in supporting the colocalization results. For example, all 9 microglial colocalized effects in AD are also found in the TWAS; when the lead SNP (and those in strong LD) are removed, the TWAS analysis is still positive, suggesting that there may be additional significant associations beyond the lead haplotype in those susceptibility loci.

In addition to neuropsychiatric diseases, we also performed TWAS of brain volumes and educational attainment (**Supplementary Figure 14 and 15 and Supplementary Table 13**). Interestingly, inhibitory neurons had the most frequent associations with the volume of subcortical brain structures (accumbens, brainstem, caudate, and putamen).

## Discussion

In this study, we constructed a single-cell transcriptomic atlas of the human neocortex by applying single-nucleus RNAseq to DLPFC samples from 424 aging individuals. Here, we focused our efforts on mapping the effect of genetic variation on gene expression: we have catalogued the genetic regulation of gene expression per brain cell types and their subtypes. We identified a large number of *cis*-eQTLs at the level of cell types and cell subtypes; this provides a significant advance over the prior existing effort, which identified only a third of the eGenes we report at the cell type level, had a much smaller sample size, collated data from different brain regions, and only evaluated *cis*-eQTLs at the cell type level. One of our principal findings is that many eGenes are only discovered when analyses are conducted at the cell subtype level (Figure 1G), highlighting an important strategic choice for future study designs: sequencing a larger number of nuclei per participant may be more important than increasing sample size.

Single nucleus analysis enabled us to detect numerous eQTL that were not detected by an eQTL analysis of bulk brain tissues, despite analyzing less than half the number of specimens used in the bulk analysis from the same ROSMAP cohorts. The comparison with the earlier snucRNAseq-based study shows that these single nucleus-derived results are replicable. Why does single nucleus eQTL analysis have enhanced power to detect eQTL? A possible explanation is cell type-and subtype-specific regulation of gene expression, consistent with the epigenomic enrichment that we have noted (Figure 4C). As we exemplified with *APOE* expression in microglia, we revealed that eGenes were often expressed in multiple brain cell types, but much of the genetic regulation is fine-tuned, probably affecting more context-specific enhancer elements. Such cell type-specific effects will be “diluted” in bulk tissue RNA profiles and thus were difficult to detect in earlier brain *cis*-eQTL maps (*5*). Here, this microglial-specific *APOE cis*-eQTL is likely to be driven by a variant in a microglial enhancer some distance away from the *APOE* gene that is brought into contact with APOE by a chromatin loop; interestingly, this variant also influences the accumulation of CAA (an amyloid-driven vasculopathy) but not parenchymal amyloid plaques or tau tangles. On the other hand, the well-known APOEε4 haplotype influences all 3 measures. This provides an important mechanistic insight: that APOE RNA expression levels secreted by microglia play a causal role in CAA while they do not affect parenchymal amyloid proteinopathy or taupathy. Microglia are certainly involved in both AD and CAA processes, but only the coding variants that define the AD susceptibility haplotype and are not strongly related to gene expression influence AD pathology. This result is consistent with existing literature reporting increased APOE levels at the site of CAA in humans (*35*) as well as data in murine models treated with an anti-APOE antibody which led to reduced CAA accumulation (*36*).

Mapping eQTL of rare cell types, such as microglia, has been a technical challenge. Recent efforts to map microglia eQTL have relied on dissociation and purification of microglia (*10*, *11*, *37*). However, enzymatic dissociation of brain tissues can induce artifactual gene expression changes, especially in microglia (*38*). Collecting and processing hundreds of fresh clinical specimens also pose logistic difficulties; these experimental concerns are alleviated when using banked, frozen samples. On the other hand, our single nucleus eQTL analysis suffered from low detection power in less common cell types, including microglia and their subtypes. This stems from such cell types receiving fewer sequencing reads, and thus fewer genes are detected and tested for eQTL; further, fewer nuclei are assigned to each subtype, yielding a noisier pseudobulk RNA expression estimate for subtypes, which will tend to reduce statistical power. One solution to this issue would be to sequence individuals more deeply by single nucleus analysis. As library preparation costs drop, this approach may become more feasible than cell purification, which is arduous for most neocortical cell types and could lead to exclusion of certain subtypes. Sample multiplexing approaches can substantially reduce costs and batch effects, as we demonstrate in this study which relied only on genetic demultiplexing without antibody-based tags, which minimizes manipulation of the samples (vs. nuclear hashing) and reduces the quantity of sequencing needed.

In reviewing results at the cell subtype level, cell subtypes generally had fewer eGenes than their parent cell type. However, subtype analysis was more productive than cell type analysis in terms of the number of eGenes per cell proportion. This seems analogous to the relationship between cell type-level eGenes and bulk eGenes. Thanks to this characteristic, subtype analysis was able to identify additional eGenes, especially in excitatory and inhibitory neurons. Deeper sequencing of individual samples and meta-analysis of subtypes would further improve detection power and are priorities for future studies.

iPSC-derived neurons and glial cell lines are an attractive model system to explore functional consequences of genetic variants related to human diseases and brain traits(*24*, *39*). Here, we confirmed that a significant fraction of our brain-derived eGenes was replicated in both iN and iAstro, despite a small sample size and thus limited statistical power for the *in vitro* model system data. Most eQTLs had the same effect direction *in vivo* and *in vitro*. However, some eQTLs, including the one affecting the *MAPT* gene, had opposite effect directions (Figure 3E and 3F), suggesting that genetic effects are highly context specific, and that caution is needed when extrapolating brain-derived genetic effects to this model system. Future epigenetic profiling of these cell lines would give insights into these context specificities and may serve as a useful resource with which to prioritize candidates for evaluation and to validate findings *in vitro*.

Genome-wide association studies have successfully identified numerous genetic variants that affect complex human traits, such as neurodegenerative diseases. However, functional follow-up of identified variants has lagged behind gene discovery efforts. Our colocalization analysis revealed cell types and even subtypes where risk variants exert their effect. This information is critical to facilitate prioritization of variants and to design functional follow-up experiments in the proper context. Furthermore, our cell type-specific TWAS identified genes whose expression are causally related to risk of neurodegenerative diseases, further expanding the list of genes to consider in functional studies. A clear result from these various disease-related analyses is that, while microglia in AD and neurons in schizophrenia, PD and educational attainment are targets of predilection for genetic susceptibility, all cell types harbor the primary effect of some disease susceptibility variants. These complex human traits – not surprisingly – are cellularly distributed, highlighting the fact that it is the sum of multiple different perturbations that lead to neurodegeneration or altered behavior: there is a systemic effect that summarizes the perturbations in individual cell types. With our results in hand, we can now begin to map out how the different cell types that we implicate in a given disease may be interacting to contribute to disease onset: we can formulate specific hypotheses to test in model systems and can narrow down functional work to the right cell type for a large number of susceptibility loci.

Overall, this manuscript reports the output of one large effort to systematically map the effect of genetic variation across a wide variety of cellular contexts in the aging brain: we use a single set of participants whose brain samples were collected prospectively using a single protocol, and all samples were processed by the same experimental and analytic pipeline. This reduces potential technical heterogeneity that can emerge when merging different sample collections and processing sites. Our use of genetic demultiplexing at large scale in single nucleus data is also useful to illustrate the robustness of our study design to reduce the number of batches without increasing sample handling. While we already help to guide the further characterization of many susceptibility loci, it is clear that many more eGenes remain to be discovered, and we highlight deeper sequencing to better resolve cell subtypes as an important aspect of the path forward.

Using a novel APOE variant that influences RNA expression in a cell-type specific manner, we also make an important insight into mechanistic differences in the deposition of two forms of amyloid, with level of APOE expression being only relevant in CAA and not in AD-related amyloid proteinopathy. This observation may have clinical relevance as the presence of CAA is a risk factor for Amyloid Related Im aging Abnormality (ARIA) following anti-amyloid antibody treatment, and APOEε4 is the major risk factor for ARIA, presumably through its effect on CAA. Thus, the new APOE variant that we report may increase risk for this important adverse event in AD immunotherapy. Finally, with its translatability to the *in vitro* system demonstrated, this resource provides an early platform for many disease-related studies as well as more fundamental studies of human genetic variation.

## Materials and Methods

### Study participants

All brain specimens were derived from two longitudinal clinical-pathological cohort studies: the Religious Orders Study (ROS), and the Memory and Aging Project (MAP) (*13*). In both cohorts, participants did not have known dementia at the time of enrollment. The participants agreed to receive clinical evaluation each year and to donate their brain at the time of death. Each study was approved by an Institutional Review Board of Rush University Medical Center. All participants signed an informed consent, Anatomic Gift Act, and repository consent. For this study, we selected 479 specimens based on availability of frozen pathologic material from the DLPFC; only participants with a post mortem interval <24 hours were considered, as in our prior studies (*40*). At the end of our data preprocessing and quality control analyses (described below), we also eliminated 19 subjects without whole genome sequence data, leaving 424 participants retained for genetic analyses. Their demographic and clinicopathologic characteristics are described in **Supplementary Table 1.**

### Whole genome sequencing

Whole genome sequencing (WGS) of ROS/MAP participants were performed as described previously (*41*). Briefly, DNA was extracted from brain or blood samples. WGS libraries were prepared using the KAPA Hyper Library Preparation Kit and sequenced on an Illumina HiSeq X sequencer as paired end reads of 150 bp. Reads were mapped to the reference human genome GRCh37 using BWA-mem, and variants were called by GATK HaplotypeCaller. For this study, the VCF file was lifted over to GRCh38 using Picard LiftoverVcf, and only variants that passed the GATK filter were used. These WGS VCF files are available in the AD Knowledge Portal (https://www.synapse.org/#!Synapse:syn11724057).

### Library preparation and sequencing of single nuclei

Each batch of samples for library construction consisted of 8 participants, except batch B63 that consisted of 7 participants. Batches were designed to balance clinical and pathological diagnosis and sex as much as possible (**Supplementary Figure 1**). Dorsolateral Prefrontal Cortex (DLFPC) tissue specimens were received frozen from the Rush Alzheimer’s Disease Center. We observed variability in the morphology of these tissue specimens with differing amounts of gray and white matter and presence of attached meninges. Working on ice throughout, we carefully dissected to remove white matter and meninges, when present. The following steps were also conducted on ice: about 50-100mg of gray matter tissue was transferred into the dounce homogenizer (Sigma Cat No: D8938) with 2mL of NP40 Lysis Buffer [0.1% NP40, 10mM Tris, 146mM NaCl, 1mM CaCl_2_, 21mM MgCl_2_, 40U/mL of RNAse inhibitor (Takara: 2313B)]. Tissue was gently dounced while on ice 25 times with Pestle A followed by 25 times with Pestle B, then transferred to a 15mL conical tube. 3mL of PBS + 0.01% BSA (NEB B9000S) and 40U/mL of RNAse inhibitor were added for a final volume of 5mL and then immediately centrifuged with a swing bucket rotor at 500g for 5 mins at 4°C. Samples were processed 2 at a time, the supernatant was removed, and the pellets were set on ice to rest while processing the remaining tissues to complete a batch of 8 samples. The nuclei pellets were then resuspended in 500ml of PBS + 0.01% BSA and 40U/mL of RNAse inhibitor. Nuclei were filtered through 20um pre-separation filters (Miltenyi: 130-101-812) and counted using the Nexcelom Cellometer Vision and a 2.5ug/ul

DAPI stain at 1:1 dilution with cellometer cell counting chamber (Nexcelom CHT4-SD100-002). 5,000 nuclei from each of 8 participants were then pooled into one sample, and the 40,000 nuclei in around 15-30ul volume were loaded into two channels on the 10X Single Cell RNA-Seq Platform using the Chromium Single Cell 3’ Reagent Kits version 3. Libraries were made following the manufacturer’s protocol. Briefly, single nuclei were partitioned into nanoliter scale Gel Bead-In-Emulsion (GEMs) in the Chromium controller instrument where cDNA share a common 10X barcode from the bead. Amplified cDNA is measured by Qubit HS DNA assay (Thermo Fisher Scientific: Q32851) and quality assessed by BioAnalyzer (Agilent: 5067-4626). This WTA (whole transcriptome amplified) material was diluted to <8ng/ml and processed through v3 library construction, and resulting libraries were quantified again by Qubit and BioAnalzyer. Libraries from 4 channels were pooled and sequenced on 1 lane of Illumina HiSeqX by The Broad Institute’s Genomics Platform, for a target coverage of around 1 million reads per channel. The same libraries of batches B10–B63 were resequenced at The New York Genome Center using Illumina NovaSeq 6000. Sequencing data of both Broad Institute and New York Genome Center were used for analysis.

### Processing of single-nucleus RNAseq reads

For each batch of snRNAseq FASTQ files, the CellRanger software (v6.0.0; 10x Genomics) was used to map reads onto the reference human genome GRCh38, to collapse reads by UMI, and to count UMI per gene per droplet. As a transcriptome model, the “GRCh38-2020-A” file set distributed by 10X Genomics was used. The “--include-introns” option was set to incorporate reads mapped to intronic region of nuclear pre-mRNA into UMI counts. To call cells among the entire droplets, the “remove-background” module of CellBender (https://github.com/broadinstitute/CellBender) was applied to raw UMI count matrices. The admixture of ambient RNA was estimated and subtracted from UMI counts by CellBender. These filtered UMI count matrices were used in the subsequent analyses. All raw and processed data are available through the AD Knowledge Portal (https://www.synapse.org/#!Synapse:syn31512863).

### Demultiplexing

Because our snRNAseq library consisted of nuclei from eight individuals, each nucleus was assigned back to its participant of origin using each nucleus’ genotype data obtained from the snRNAseq reads. We employed two different procedures, depending on whether all eight individuals had been genotyped with WGS. When eight individuals were genotyped, we used demuxlet software (*17*). From the WGS-based VCF file of 1,196 ROS/MAP individuals, we extracted SNPs that were in transcribed regions, passed a filter of GATK, and at least one of the eight individuals had its alternate allele. The extracted SNP genotype data were fed to demuxlet along with BAM file generated by CellRanger. When less than eight individuals were genotyped, we used freemuxlet (https://github.com/statgen/popscle), which clusters droplets based on SNPs in snRNAseq reads and generates a VCF file of snRNAseq-based genotypes of the clusters. The number of clusters was specified to be eight. The snRNAseq-based VCF file was filtered for genotype quality > 30 and compared with available WGS genotypes using the bcftools gtcheck command. Each WGS-genotyped individual was assigned to one of droplet clusters by visually inspecting a heatmap of the number of discordant SNP sites between snRNAseq and WGS. The above two procedures converged to a table that mapped droplet barcodes onto inferred individuals. Each BAM file generated by CellRanger was split into eight per-individual BAM files, each of which contained reads from distinct individual, using subset-bam (https://github.com/10XGenomics/subset-bam). UMI count matrices filtered by CellBender were split into eight per-individual UMI count matrices.

### Quality control

Among 479 specimens analyzed by snRNAseq, 19 specimens were excluded from our analyses because they did not have WGS genotypes. Also, three specimens were excluded at the stage of freemuxlet-based demultiplexing because they had ambiguity in assignment of droplet clusters to individual genotypes. To identify and exclude potential sample swaps in the remaining 457 specimens, we assessed concordance of genotypes between snRNAseq and WGS. LOD scores, a metric of genotype concordance, were computed by comparing the per-individual BAM files with WGS genotypes of matched individuals using Picard CrosscheckFingerprints (ver. 2.25.4). We used a haplotype map downloaded from https://github.com/naumanjaved/fingerprint_maps. After inspecting a histogram of LOD scores, ten specimens whose LOD scores were less than 50.0 were filtered out. These specimens received few cells by the demultiplexing procedure and were set aside from future reprocessing. As another measure to detect sample swaps, we checked RNA expression levels of *XIST* gene and confirmed that they were consistent with clinical sex of the remaining 447 specimens. Three individuals were further excluded because they failed quality control of WGS; two were marked as potential sample swaps among WGS, and the other was marked as an outlier based on genotype principal component analysis. The latter individual was discarded given the preference to have a genetically homogeneous set of individuals for QTL mapping.

Four specimen-level sequencing metrics were computed from the per-specimen UMI count matrices: estimated number of cells, median UMI counts per cell, median genes per cell, and total genes detected. After inspecting these metrics, eight specimens whose median UMI counts per cell were less than 1,500 were excluded. Among the remaining 436 specimens, 12 individuals were found to be sequenced twice in distinct batches. After comparing sequencing metrics, one of these duplicates were excluded from further analyses. After these quality control processes and matching to whole genome sequence data was available, 424 individuals remained.

### Cell type classifications

Based on the cell-type annotations of our prior work (*16*), we fitted a regularized logistic-regression classifier and calculated cell-type probabilities for all 127 single-nucleus libraries. Then, low-quality nuclei were determined using cell-type specific thresholds of total number of UMIs and number of unique features, and low-quality nuclei were removed. Doublets were detected by running DoubletFinder (*42*) and using the demultiplexing doublets as a partial ground-truth to define a library-specific doublets threshold. Thresholds for doublet removal was chosen to be the proportion of artificial nearest neighbors maximizing Matthew’s correlation coefficient.

Next, nuclei were split into cell-type objects. For each cell-type the gene expression matrix was normalized and scaled using Seurat’s SCTransform method (variable.features.n=4000) (*43*) followed by PCA dimensionality reduction (using RunPCA method, npcs=50, Seurat package) and the community-detection based clustering leiden algorithm (using FindClusters method, algorithm=4, Seurat package). Clustering resolution was determined by subsequent clusters expressing a distinct set of differentially expressed genes, having a distinct set of pathways, or containing nuclei originating from a specific set of donors.

### Pseudobulk expression quantification

As we mentioned, our snRNAseq libraries were prepared in duplicate. UMI counts of the two replicates from the same individual were aggregated together. For each cell type, individuals who had less than 10 cells were excluded from expression quantification for that cell type. Rare cell types were excluded from subsequent analysis if less than 10 individuals were available for expression quantification. We generated a pseudobulk UMI count matrix for each cell type by extracting UMI counts of the cell type and by aggregating the counts per gene per individual. Low expression genes were filtered out by using filterByExpr function of edgeR (version 3.30.3) with its default parameters. The pseudobulk counts were normalized by using the trimmed mean of M-values (TMM) method of edgeR, and log_2_ of counts per million mapped reads (CPM) were computed using the voom function of limma (version 3.44.3). Low expression genes whose log_2_CPM were less than 2.0 were filtered out. Batch effects were corrected using ComBat. Expression levels were quantile normalized. Pseudobulk expression of cell subtypes were quantified by the same method.

### Mapping of *cis*-eQTL

We used the Matrix eQTL version 2.3 method (*18*) to identify *cis*-eQTLs with 1 Mb of the transcription start site of each measured gene (gene expression derived using log_2_CPM). All bi-allelic single nucleotide polymorphisms (SNPs) with a minor allele frequency >0.05, a call rate >95%, and Hardy-Weinberg *p* > 10^-6^ were retained for analysis. We used a linear model for gene expression whose explanatory variables were allele counts of SNP and several covariates. Statistical significance was computed from *t*-statistic. Genotype principal components (PC) were calculated using PLINK version v1.90 (*44*), and we included the top 3 genotype PCs to account for residual population structure among these individuals of European ancestry. We also calculated expression PCs based on the RNA expression data within each cell type to identify the number of expression PCs that optimizes *cis*-eQTL discovery by regressing out non-genetic structure in the data; results of these evaluations are shown in **Supplementary Figure 14.** While there was some variation in the optimal number of PCs to include in each cell type, differences were small, and we opted to be consistent and to include the top 30 expression PCs as covariates. We also examined postmortem interval and clinical traits of individuals as covariates, but they had little impact on the number of eQTL detected (**Supplementary Figure 15**). The final set of covariates were top 3 genotype PCs, top 30 expression PCs, age, sex, post-mortem interval, study (ROS or MAP), and total number of genes detected in each participant. Multiple hypothesis correction was performed using a two-step method. Gene-wise *p*-values were computed by applying Bonferroni correction to the smallest nominal *p*-value of each gene with the number of tested SNPs for the gene. The threshold for statistical significance of eGenes was set to the false discovery rate (FDR) < 5%, where FDR was computed from gene-wise *p*-values using the Benjamini-Hochberg method. Statistical significance of eSNPs were judged by nominal *p*-values, and its threshold was set to the largest nominal *p*-value of gene-SNP pair that had FDR < 5%. Lead eSNP was selected to have the smallest *p*-value for each eGene.

### Chromatin states

Fifteen-class chromatin states of a bulk DLPFC tissue (E073) were downloaded from the FTP site of the Roadmap Epigenome project. Cell type-specific enhancers and promoters of brain cells were downloaded from USCS genome browser (https://genome.ucsc.edu/). The SNPs tested for eQTL were categorized into four groups based on whether they were eSNPs and whether they were in the chromatin state of interest. A contingency table was constructed by the number of SNPs in the four categories. Log odds ratio and *p*-values of Fisher’s exact test were computed using R package “epitools”.

### Bulk RNA-seq and eQTL

DLPFC tissues of postmortem brains from 1,092 ROSMAP participants were profiled using bulk RNA-seq. Transcripts per million (TPM) values of gene expression levels were computed and normalized using the TMM method. Outlier samples were removed based on gene expression profiles. Low-expression genes with a median TPM less than 10 were excluded. The normalized TPM values were log_2_-transformed and adjusted for age, sex, batch, and technical confounding factors using a linear regression. As technical confounding factors, we used library size, percentage of coding bases, percentage of aligned reads, percentage of ribosomal bases, percentage of UTR base, median 5 prime to 3 prime base, median CV coverage, pmi, and study index of ROS or MAP. A similar pipeline as for cell type specific eQTL was applied to estimate bulk eQTLs. Analogous threshold criteria were used for declaring significance. We compared the cell type specific eQTLs against the bulk eQTLs in a couple of ways. First, we examined which significant cell type specific eQTLs are distinct from the bulk eQTLs as well as which overlap. Second, we used π_1_ statistics to estimate the proportion of cell type specific eQTLs that also display associations in bulk. Specifically, restricting to gene-SNP pairs that passed significance threshold of each cell type, we extracted nominal *p*-values of the same gene-SNP pairs from bulk eQTL and estimated π_1_ using the “pi0est” function of “qvalue” R package. We also estimated π_1_ in the reverse direction to examine the proportion of bulk eQTLs displaying associations in cell type specific eQTLs.

### Meta-PheWAS

We performed a combined phenome-wide association analysis using eMERGE-III and UK Biobank (UKBB) datasets (*32*, *33*). Details on genotyping and quality control analyses for eMERGE-III and UK Biobank datasets have been described previously (*45*–*47*). The 102,138 genotyped eMERGE-III participants had a total of 20,783 ICD-9 codes mapped to 1,817 distinct phecodes. The 460,363 UK Biobank participants with genotype data had a total of 10,221 ICD-9 codes mapping to 1,817 phecodes. Phenome-wide association analyses were performed using the PheWAS R package 0.99.5.5 (*48*). The package uses pre-defined “control” groups for each phecode “case” grouping and two phecode occurrences defined as a case. In total, all 1,817 phecodes were tested across both datasets using logistic regression with the phecode case-control status as the outcome and SNP genotype, sex, age, site, and 5 principal components of ancestry as predictors. We then combined the analysis across both cohorts using the Meta-PheWAS approach under a fixed effects model, as implemented in the PheWAS R package. To declare phenome-wide significance, we set the Bonferroni corrected statistical significance threshold at 2.75 x 10^-5^ corrected for 1,817 phecodes tested.

### Induced neurons and astrocytes

iPSC lines were generated using Sendai virus-mediated reprogramming from deceased ROSMAP participants for whom neuropathological traits were measured. The generation and characterization of these lines has been described previously (*24*). iPSC lines were differentiated to excitatory neuronal fates using a NGN2 protocol, and bulk RNA-seq performed at day 21 as described previously (*24*). Briefly, iPSCs were plated in mTeSR1 media at a density of 95K cells/cm^2^ on Matrigel coated plates for viral transduction. Media was changed from StemFlex to mTeSR1 as we found better transduction viability with mTeSR1. Viral plasmids were obtained from Addgene (plasmids #19780, 52047, 30130). FUdeltaGW-rtTA was a gift from Konrad Hochedlinger (Addgene plasmid # 19780). Tet-O-FUW-EGFP was a gift from Marius Wernig (Addgene plasmid # 30130). pTet-ONgn2-puro was a gift from Marius Wernig (Addgene plasmid # 52047). Lentiviruses were obtained from Alstem with ultrahigh titers (10^9^) and used at the following concentrations: pTet-O-NGN2-puro: 0.1 ul/ 50K cells; Tet-O-FUW-eGFP: 0.05ul/ 50K cells; Fudelta GW-rtTA: 0.11ul/50K cells. Transduced cells were dissociated with Accutase and plated onto Matrigel-coated plates at 50,000 cells/cm^2^ in mTeSR1 (day 0). On day 1, media was changed to KSR media with doxycycline (2 ug/ml, Sigma). Doxycyline was maintained in the media for the remainder of the differentiation. On day 2, media was changed to 1:1 KSR: N2B media with puromycin (5 ug/ml, Gibco). Puromycin was maintained in the media throughout the differentiation. On day 3, media was changed to N2B media + 1:100 B27 supplement (Life Technologies), and puromycin (10 ug/ml). From day 4 on, cells were cultured in NBM media + 1:50 B27 + BDNF, GDNF, CNTF (10 ng/ml, Peprotech). Formulation of induced neuron protocol media is as follows. KSR media: Knockout DMEM, 15% KOSR, 1x MEM-NEAA, 55 uM beta-mercaptoethanol, and 1x GlutaMAX (Life Technologies). N2B media: DMEM/F12, 1x GlutaMAX (Life Technologies), 1x N2 supplement B (Stemcell Technologies), and 0.3% dextrose (D-(+)-glucose, Sigma). NBM media: Neurobasal medium, 0.5x MEM-NEAA, 1x GlutaMAX (Life Technologies), and 0.3% dextrose (D-(+)-glucose, Sigma).

iPSC-derived astrocytes (iAstros) were differentiated following a previously published paper (*49*) with minor modifications (*24*). iAstros were collected for bulk RNA-seq at day 28 of differentiation. In short, iPSCs were plated at 95k cells/cm^2^ on a growth factor reduced matrigel (Corning #354230) coated plate, then were transduced with three lentiviruses – TetO Sox9 puro (Addgene plasmid #117269), TetO Nfib Hygro (Addgene plasmid #117271), and FUdeltaGW-rtTA (Addgene plasmid #19780). The cells were then dissociated with Accutase, plated at 200,000 cells/cm^2^ using Stemflex and ROCK inhibitor (10uM) (D0). From D1 to D6, the media was gradually switched from Expansion media (DMEM/F12, 10% FBS, 1% N2 supplement, 1% Glutamax) to FGF media (Neurobasal media, 2% B27, 1% NEAA, 1% Glutamax, 1% FBS, 8ng/ml FGF, 5ng/ml CNTF, 10ng/ml BMP4). On day 8, cells are replated at 84k cells/cm^2^ and from D8 to the end of differentiation D21, cells were cultured with Maturation media (1:1 DMEM/F12 and Neurobasal, 1% N2, 1% Glutamax, 1% Sodium Pyruvate, 5ug/ml N-acetyl cysteine, 5ng/ml heparin-binding EGF-like GF, 10ng/ml CNTF, 10ng/ml BMP4, 500ug/ml dbcAMP) and fed every 2-3 days.

For iNs, at least 250 ngs of total RNA input was oligo(dT) purified, then double-stranded cDNA was synthesized using SuperScript III Reverse Transcriptase with random hexamers. Sequencing libraries were generated by processing 1 ng of generated cDNA through the Illumina Nextera Tagmentation library protocol. RNA and cDNA were quantified on a TapeStation 4200. Multiplexed libraries were sequenced on an Illumina NextSeq 500 to an average depth of 24 million mapped paired-end reads (150 bases) per sample. RNAseq reads were quality trimmed, then quantified using the Kallisto pseudoalignment quantification program (v0.43.1) (*50*) running 100 bootstraps against a Kallisto index generated from GRCh38. Kallisto quantified samples were compared using Sleuth (*51*) (v0.30.0) in R Studio (v3.6.1 of R; v1.2.5019 of R Studio). Differentially expressed genes were identified using a Wald test after controlling for library preparation batch. Expression values were exported from the Sleuth object as normalized TPM values. The RNAseq library preparation of all ROSMAP libraries took several rounds to complete. To generate a master expression matrix the exported expression vectors were combined, quantile normalized, then batch effects were regressed using the ComBat algorithm in the SVA (v3.34.0) package in R. Any gene with very low average expression (0.6TPM for gene level analysis, and 0.2TPM for transcript level analysis) was excluded from the matrix. Samples with abnormally high levels of either *Oct2/3* or *LEFTY2* (>20TPM) were suspected to have a high percentage of undifferentiated cells and were removed from the analysis

### COLOC

The COLOC package (version 5.1.0) applied the approximate Bayes factor (ABF) colocalization hypothesis, which conducted using the functionLof *coloc.abf()*, which is under a single causal variant assumption. Under ABF analysis, the association of the trait with SNPs can be achieved by calculating the posterior probability (value from 0 to 1), with the value of 1 indicating the causal SNP. In addition, the ABF analysis has 5 hypotheses, where *PP.H0.abf* indicates there is neither an eQTL nor a GWAS signal at the loci; *PP.H1.abf* indicates the locus is only associated with the GWAS; *PP.H2.abf* indicates the locus is only associated with the eQTL; *PP.H3.abf* indicates that both the GWAS and eQTL are associated but to a different genetic variant; *PP.H4.abf* indicates that the eQTL and the GWAS are associated to the same genetic variant. With the posterior probability of each SNP and aiming to find the casual variants between the GWAS and eQTL, we focused on extracting the PP.H4 value for each SNP in our study. For AD GWAS (*30*), we used the reported lead SNPs of 38 loci. For each locus, we searched for the eSNPs that are within 500 Kb of the lead SNP, and listed eGenes that were paired with the eSNP. We then obtained the eGenes *cis*-eQTL output around the lead eSNP within 1 Mb window size. In addition, we extracted GWAS summary statistics around the reported 38 lead SNP. At last, we conducted COLOC for respective pair of eGene-eQTL and eSNP-GWAS for each cell type. Similarly, for PD GWAS (*52*), there were 90 independent genome-wide significant risk loci. We picked one SNP that has the smallest P value from each locus as the lead SNP for COLOC analysis. For schizophrenia GWAS (*53*), 270 risk loci were identified as relating to schizophrenia. The SNP from each locus was used for COLOC analysis. For ALS GWAS (*54*), 15 risk loci were identified as relating to ALS, and the SNP from each locus was used for COLOC analysis. Besides the neurodegenerative diseases, we also conducted colocalization analysis on other brain traits, i.e., educational attainment (*55*) and brain volume. In terms of brain volume, we conducted COLOC using GWAS summary statistics of intracranial volume (*56*), hippocampal volume (*57*), subcortical volume (*58*), and cortical surface area and thickness (*59*).

### TWAS

We used pseudobulk RNA-seq data and genotypes from ROSMAP (424 brain subjects) to impute the *cis* genetic component of expression into multiple GWAS summary statistics as we mentioned in the COLOC analysis. The complete TWAS pipeline is implemented in the FUSION (ver. Oct. 1, 2019) suite of tools (*60*). The details steps implemented in FUSION are as follows. First, we estimated the heritability of gene expression and stopped if not significant. We estimated using a robust version of GCTA-GREML (*61*), which generates heritability estimates per feature as well as the likelihood ratio test *P* value. Only features that have a heritability of *P*L<L0.05 were retained for TWAS analysis. Second, the expression weights were computed by modeling all *cis*-SNPs (±1LMb from the transcription start site) using best linear unbiased prediction, or modeling SNPs and effect sizes with Bayesian sparse linear mixed model, least absolute shrinkage and selection operator, Elastic Net and top SNPs (*60*, *62*). A cross-validation for each of the desired models were then performed. Third, a final estimate of weights for each of the desired models was performed and the results were stored. The imputed unit is treated as a linear model of genotypes with weights based on the correlation between SNPs and expression in the training data while accounting for linkage disequilibrium (LD) among SNPs. To account for multiple hypotheses, FDR corrected *p*-value threshold (FDR ≤ 0.05) was used to define significant TWAS associations. snucRNA-sequencing from each cell subtypes (sample size ≥ 100) was also imputed for TWAS analysis.

## Acknowledgments

We thank all the participants of ROS/MAP study for their participation and generous donation of brains.

## Funding

This work was funded by NIH grants U01AG061356 (De Jager/Bennett), RF1AG057473 (De Jager/Bennett), and U01AG046152 (De Jager/Bennett) as part of the AMP-AD consortium, as well as NIH grants R01AG066831 (Menon), K25DK128563 (Khan), and U01AG072572 (De Jager/St George-Hyslop).

### Author contributions

Conceptualization: V.M., P.L.D.

Investigation: M.F., Z.G., L.Z., C.M., C.C.W., B.N., G.S.G., O.R.-R., D.P., L.A.-Z., H.L.,

R.V.P., A.K., B.N.V., K.K., C.J.Y, H.-U.K., G.W., A.R., N.H., J.A.S., Y.W., T.Y.-P., S.M., D.A.B., V.M., P.L.D.

Funding acquisition: D.A.B., V.M., P.L.D. Supervision: V.M., P.L.D.

Writing – original draft: M.F., Z.G., L.Z., P.L.D.

Writing – review & editing: M.F., C.C.W., B.N., G.S.G., K.K., C.J.Y, H.-U.K., T.Y.-P., D.A.B., P.L.D.

### Competing interests

A.R. is a co-founder and equity holder of Celsius Therapeutics, is an equity holder in Immunitas, and was a scientific advisory board member of ThermoFisher Scientific, Syros Pharmaceuticals, Neogene Therapeutics, and Asimov until 31 July 2020. Since 1 August 2020, A.R. is an employee of Genentech with equity in Roche. O.R.-R. has been an employee of Genentech since 19 October 2020. She has given numerous lectures on the subject of single-cell genomics to a wide variety of audiences and, in some cases, has received remuneration to cover time and costs. O.R.-R. and A.R. are co-inventors on patent applications filed at the Broad Institute of MIT and Harvard related to single-cell genomics. Since 3 May 2021, D.P. is an employee of Genentech with equity in Roche.

### Data and materials availability

Raw sequence data are available at Synapse (https://www.synapse.org/#!Synapse:syn31512863). Variant Call Format (VCF) files of WGS are available at Synapse (https://www.synapse.org/#!Synapse:syn11724057).

**Supplementary Materials**

Materials and Methods

Supplementary Figures 1 to 17

Supplementary Tables 1 to 13

