## Supplementary Figures for "Cell-subtype specific effects of genetic variation in the aging and Alzheimer cortex"

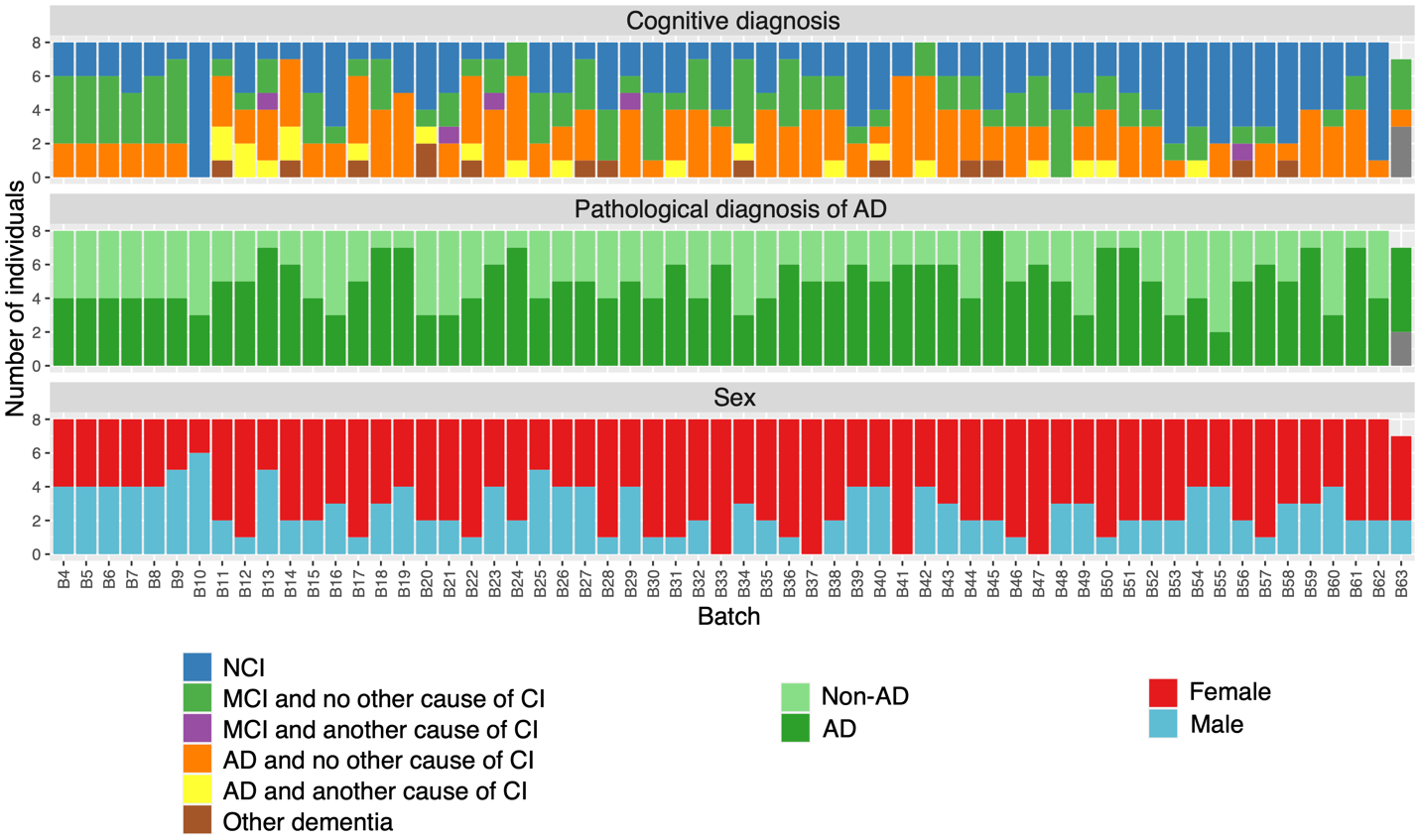

**Supplementary Figure 1. Batch design. All batches, except B63, contains nuclei from 8 individuals. NCI, no cognitive impairment; MCI, mild cognitive impairment; CI, cognitive impairment; AD, Alzheimer's disease. Gray bar shows missing data.**

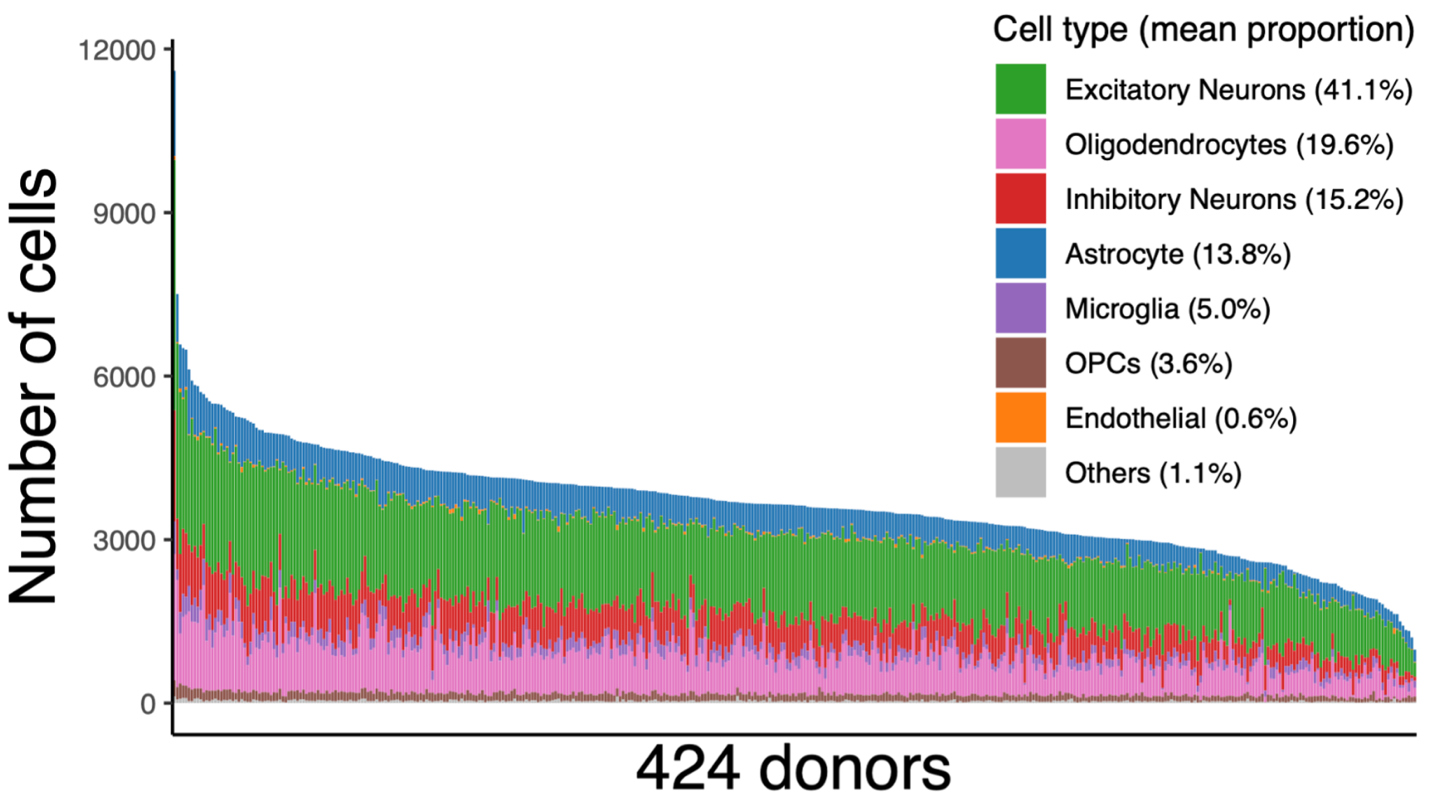

**Supplementary Figure 2. Number of cell types in 424 donors.**

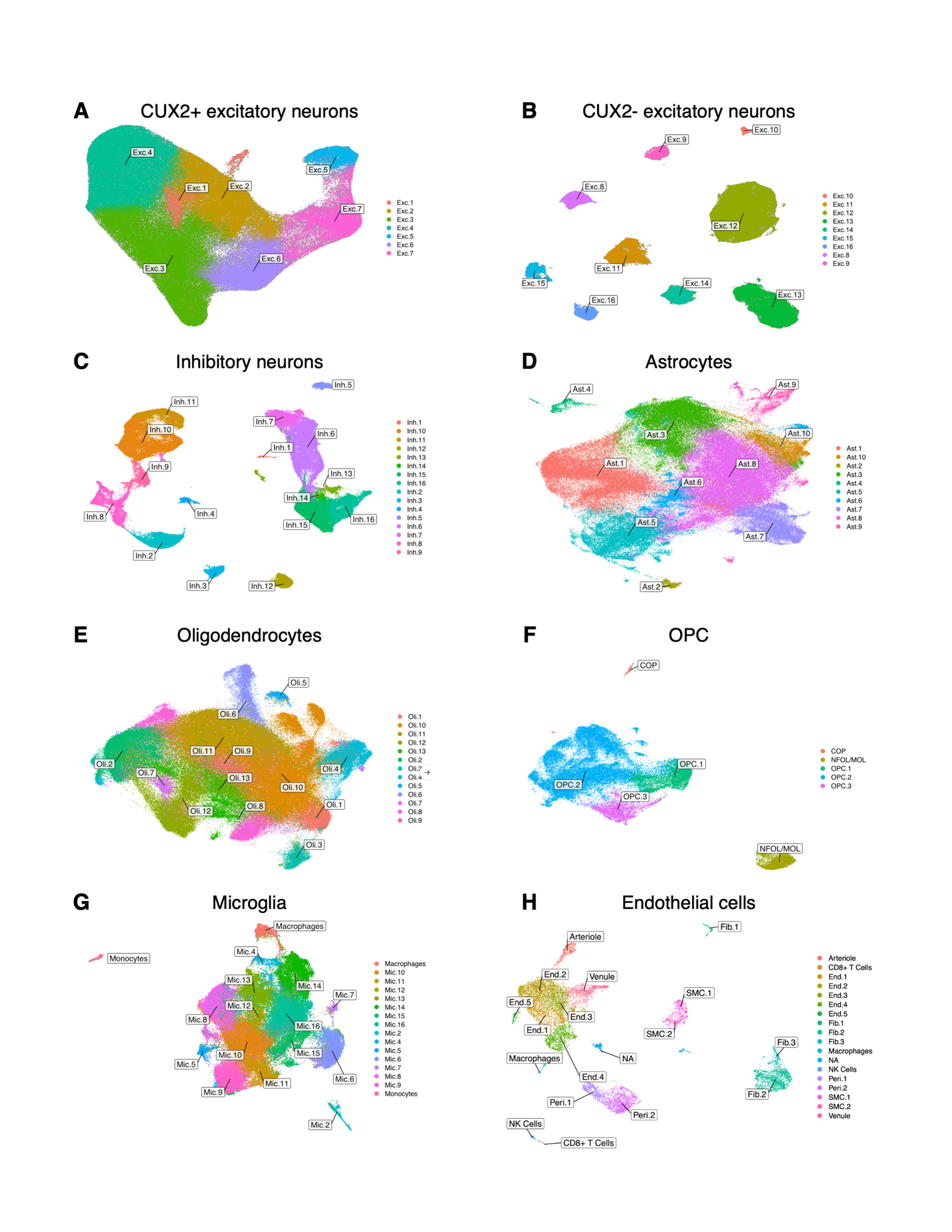

**Supplementary Figure 3. UMAP visualization of 92 cell subtypes. Macrophages appear in (G) and (H).**

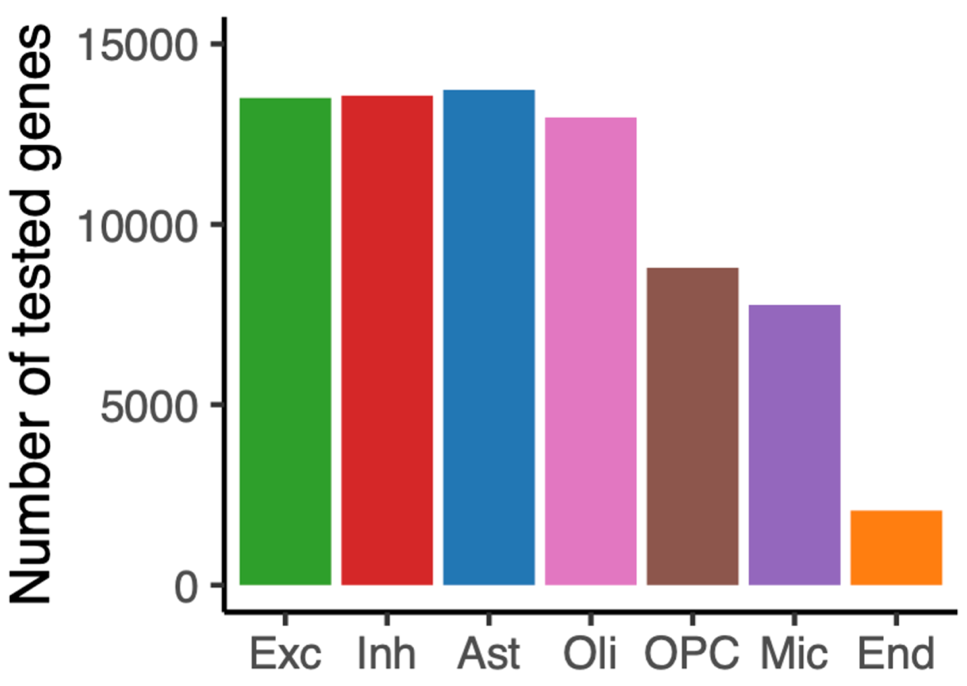

**Supplementary Figure 4. Number of genes tested for eQTL per cell type. All the cell types used the same transcriptome model “GRCh38-2020-A”, which contains 36,601 annotated genes, for gene expression quantification. However, after filtering of low expression genes, varying number of genes remained.**

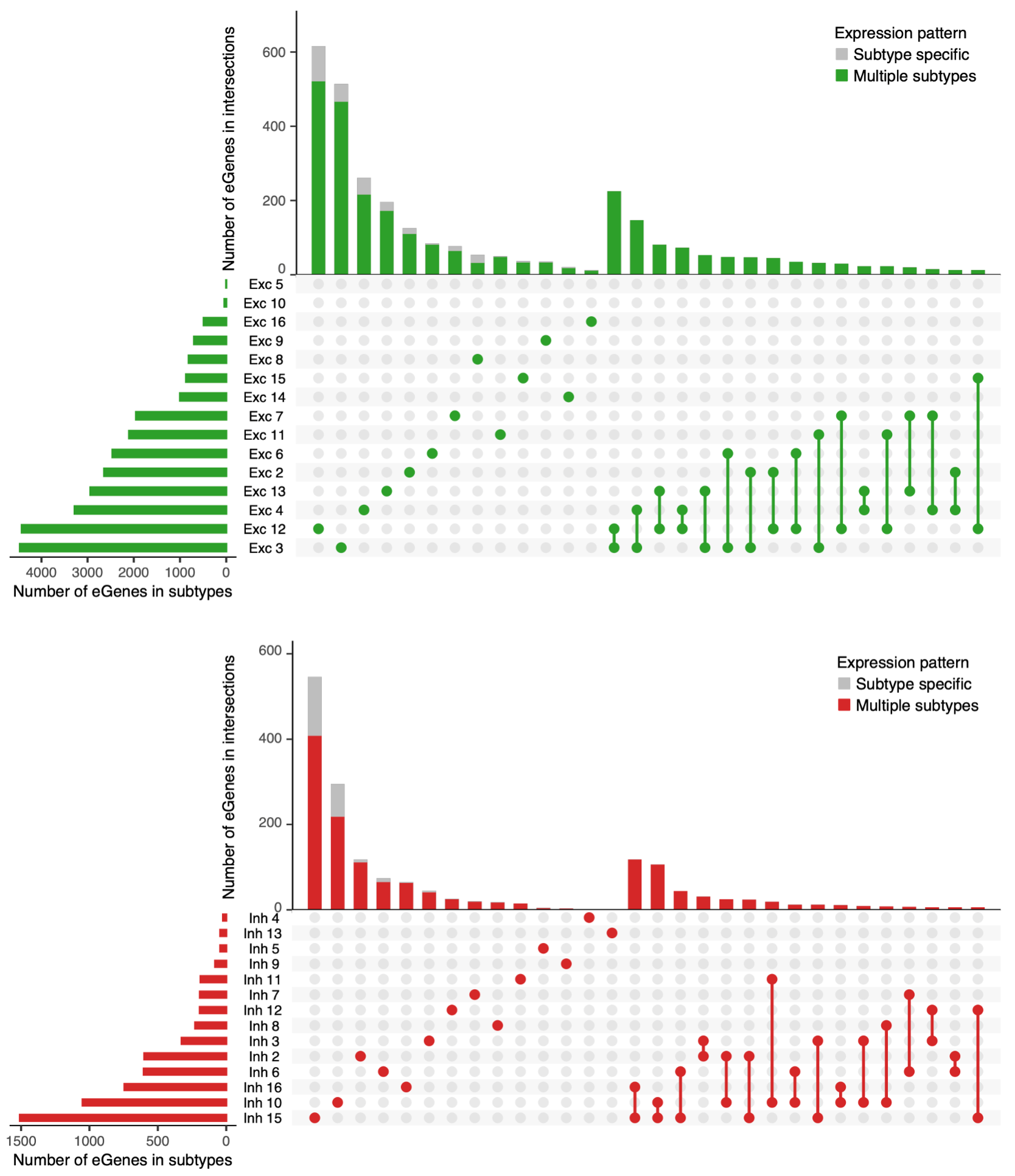

**Supplementary Figure 5. (continued)**

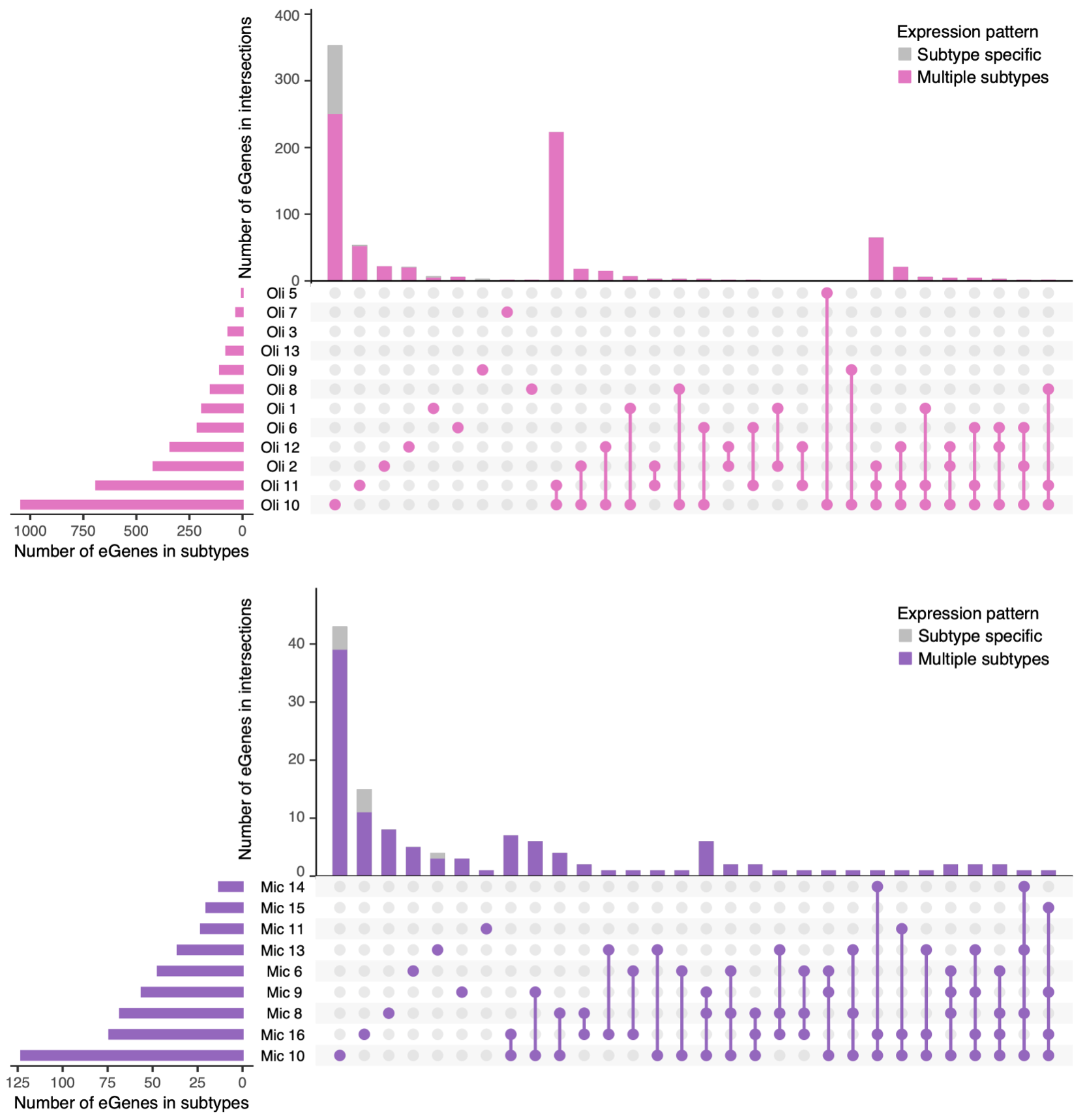

**Supplementary Figure 5. (continued)**

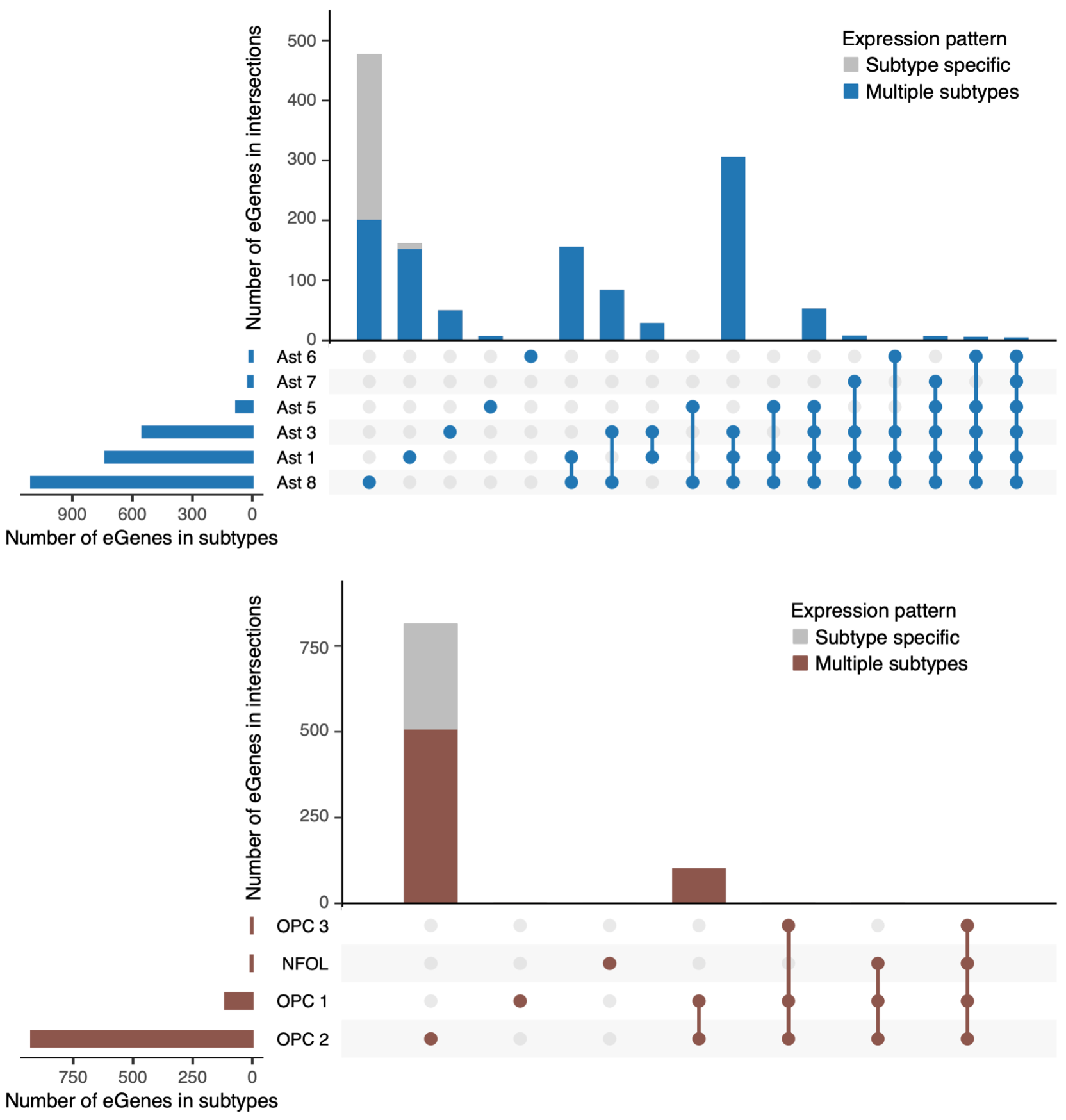

**Supplementary Figure 5. Subtype-specificity of eGenes. Number of eGenes that were unique to or shared between cell subtypes. Up to 30 intersections are shown. In the vertical bar chart, proportion of eGenes specifically expressed in one subtype are colored gray, and that expressed in two or more subtypes are colored differently.**

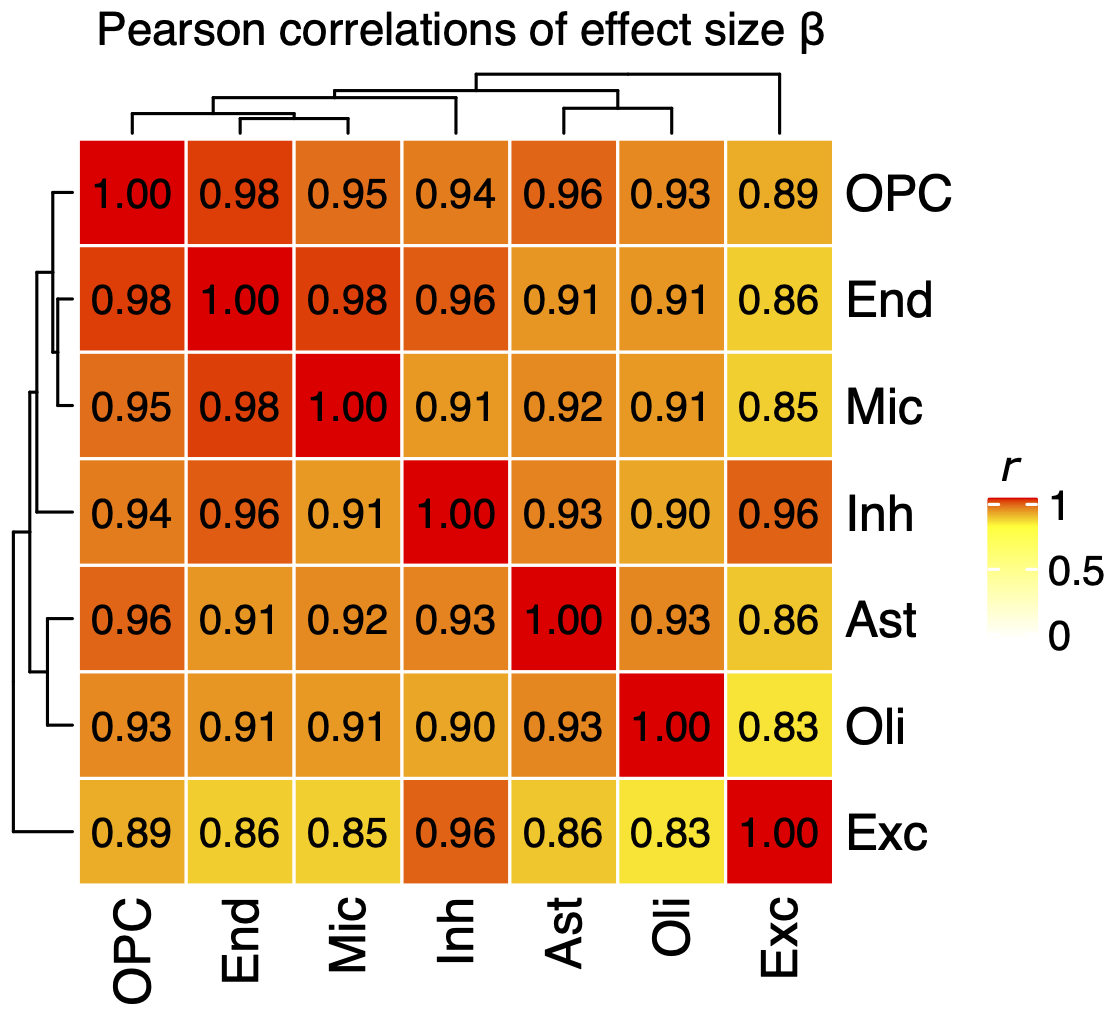

**Supplementary Figure 6. Pearson correlations of effect sizes *β* in *cis*-eQTL shared between two cell types. Rows and columns are hierarchically clustered based on the Pearson correlations.**

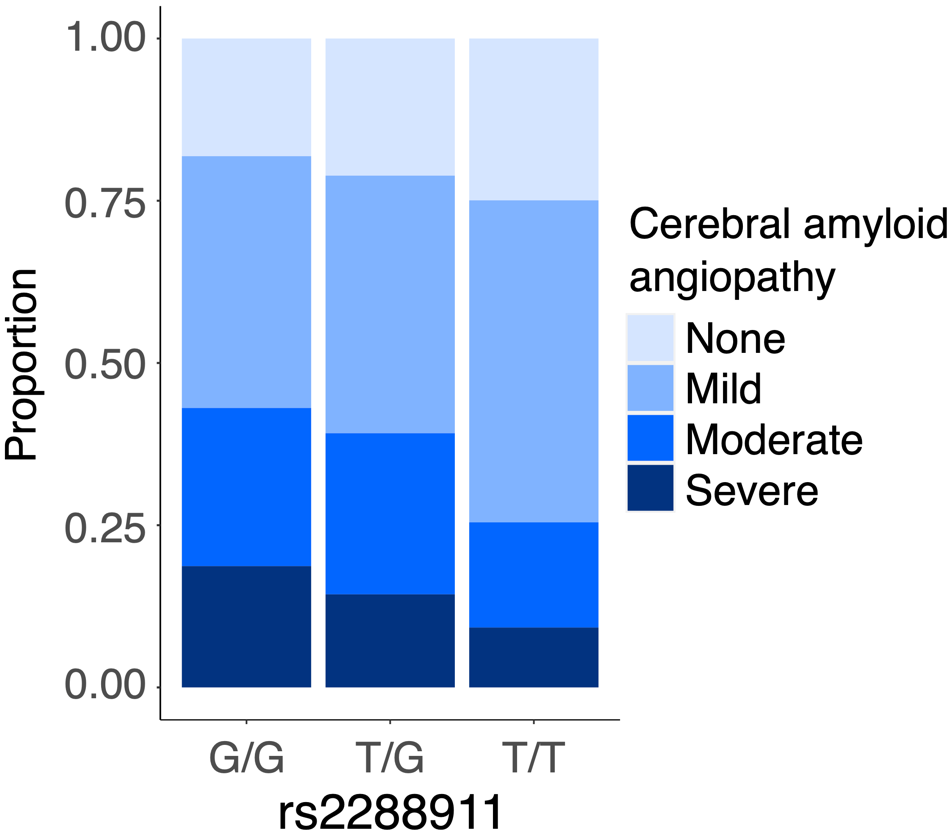

**Supplementary Figure 7. Association between rs2288911 and cerebral amyloid angiopathy (CAA) in ROSMAP cohort. The SNP rs2288911 is a microglia-specific *cis*-eQTL of *APOE*. The *y* axis shows the proportion of individuals with CAA burden per each genotype. CAA burden is a semiquantitative summary of amyloid deposition in vessels in four neocortical regions: midfrontal, midtemporal, parietal, and calcarine cortices.**

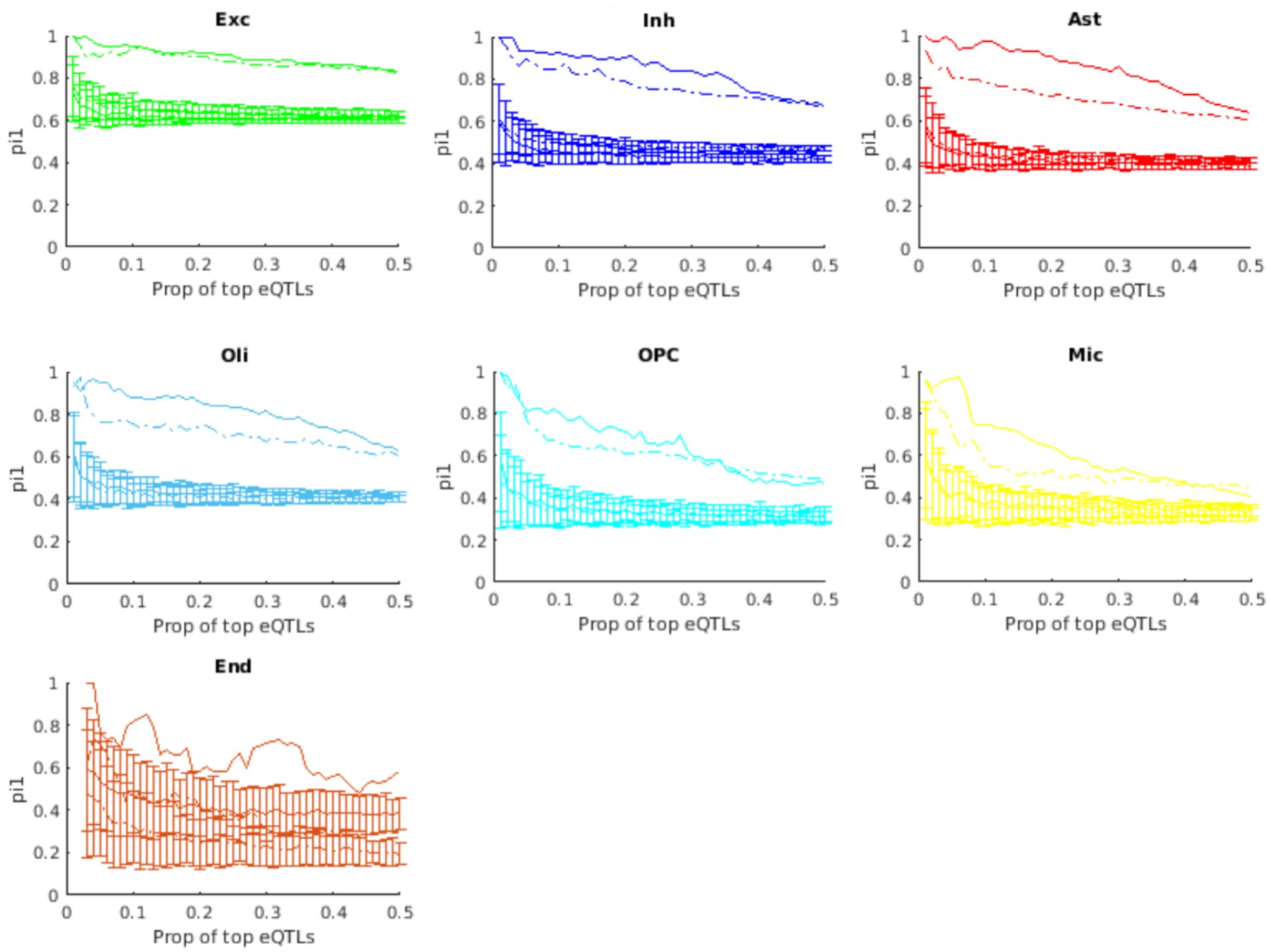

**Supplementary Figure 8. Comparison of bulk eQTL (n = 1,092) vs single-nucleus eQTL using π_1_ statistic.** **To make π_1_ statistics more independent from a choice of significance threshold, here we sorted eQTLs by *p*-values and used varying proportion (up to 50%) of top eQTLs as reference of π_1_ computation. For each cell type, four curves are drawn: Solid line without error bar, observed values of π_1_, where bulk eQTL was query. Dashed line without error bar, observed values of π_1_, where bulk eQTL was reference. Solid line with error bars, null distribution of π_1_, where bulk eQTL was query. Dashed line with error bars, null distribution of π_1_, where bulk eQTL was reference.**

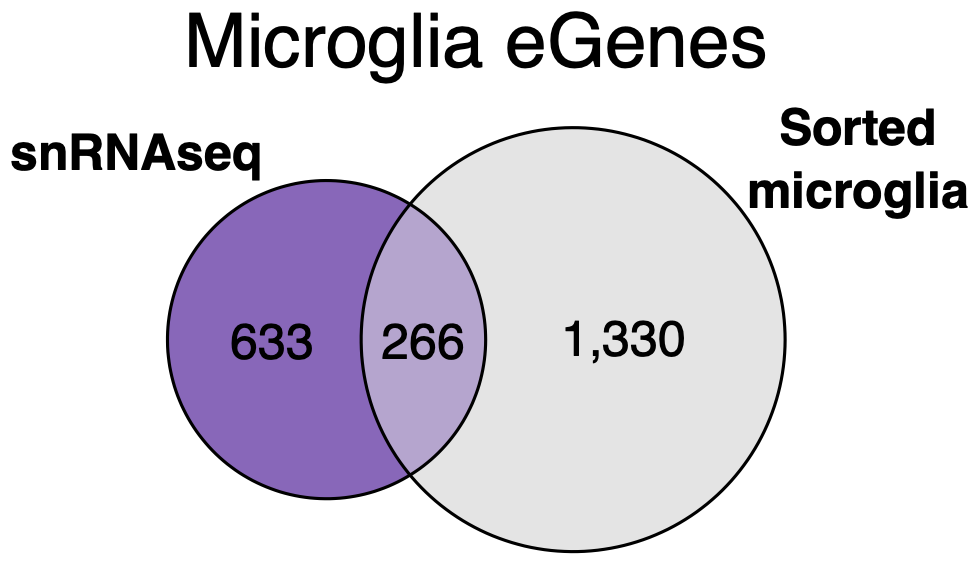

**Supplementary Figure 9. Number of microglia eGenes. Microglia eGenes detected by single-nucleus RNAseq (this study) were compared with eGenes detected by RNA-seq of microglia sorted from medial frontal gyrus (de Paiva Lopes *et al*., 2022; local false sign rate < 0.05).**

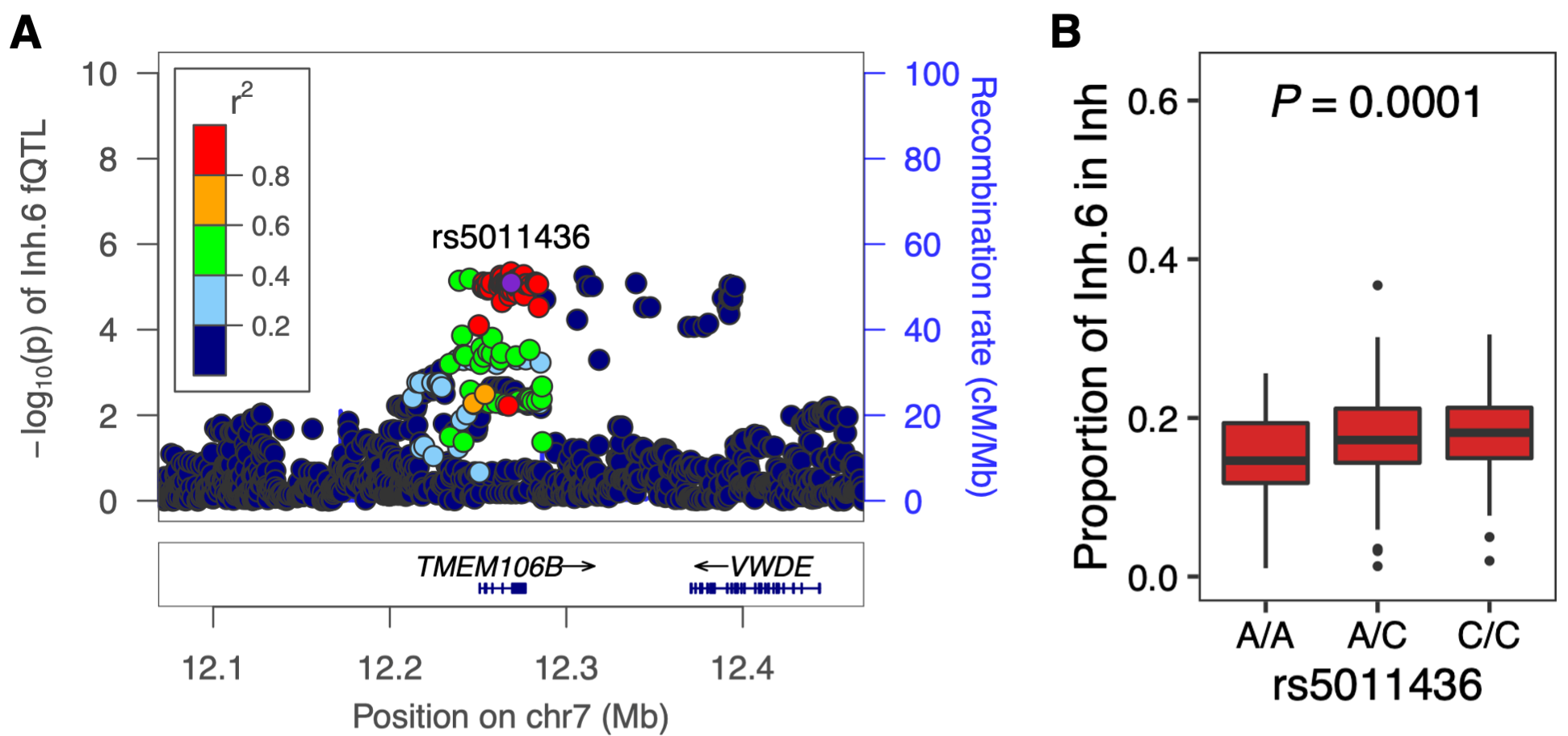

**Supplementary Figure 10. Fraction QTL of inhibitory neuron 6 (Inh.6). (A) Locuszoom plot around the *TMEM106B* locus. As a reference SNP, we used rs5011436, which is a lead risk SNP of AD in this locus according to Wightman *et al*., 2021. The *y*-axis shows -log_10_ of *p*-values for Inh.6 fQTL (B) Genotypes of rs5011436 and proportion of Inh.6.**

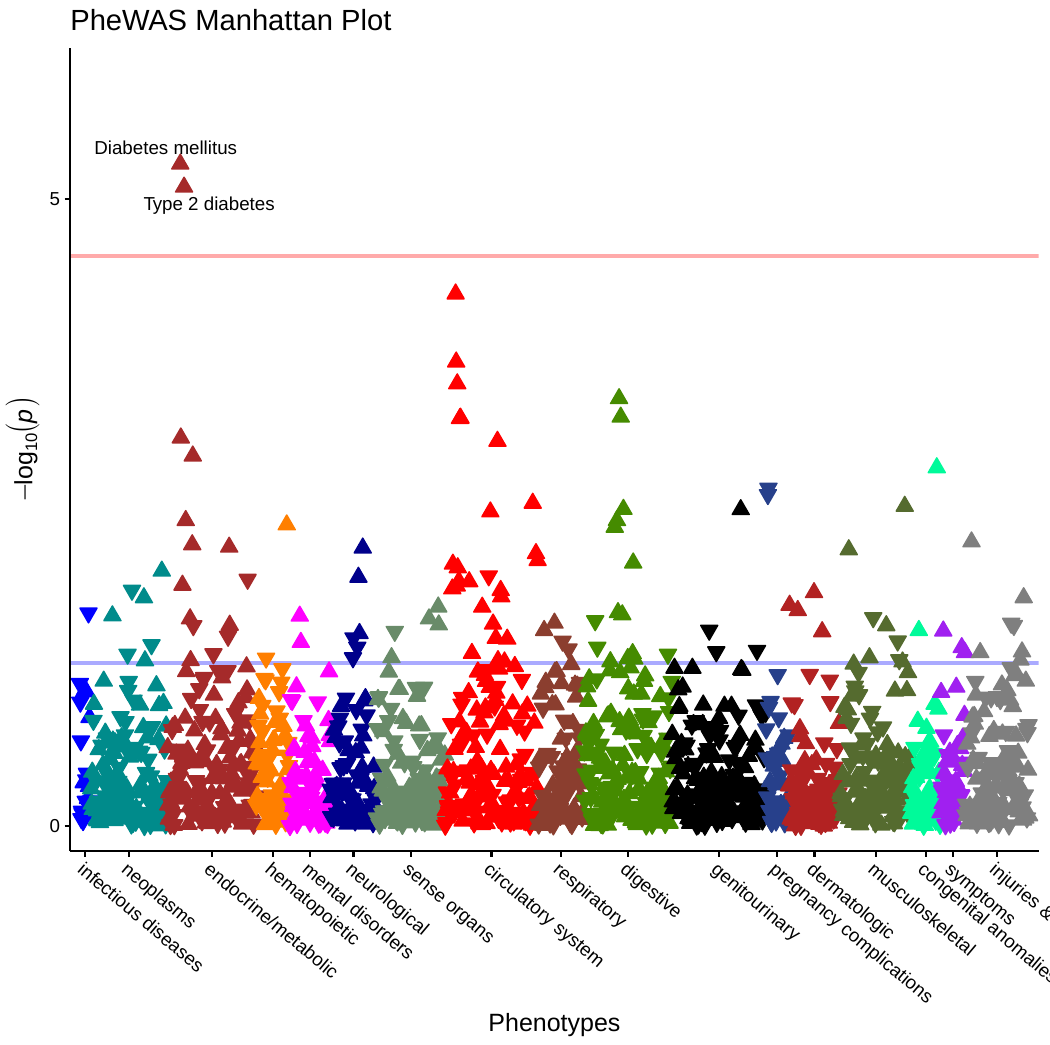

**Supplementary Figure 11. Manhattan plot of meta-PheWAS for SNP rs5011436 in UK Biobank and eMERGE3.**

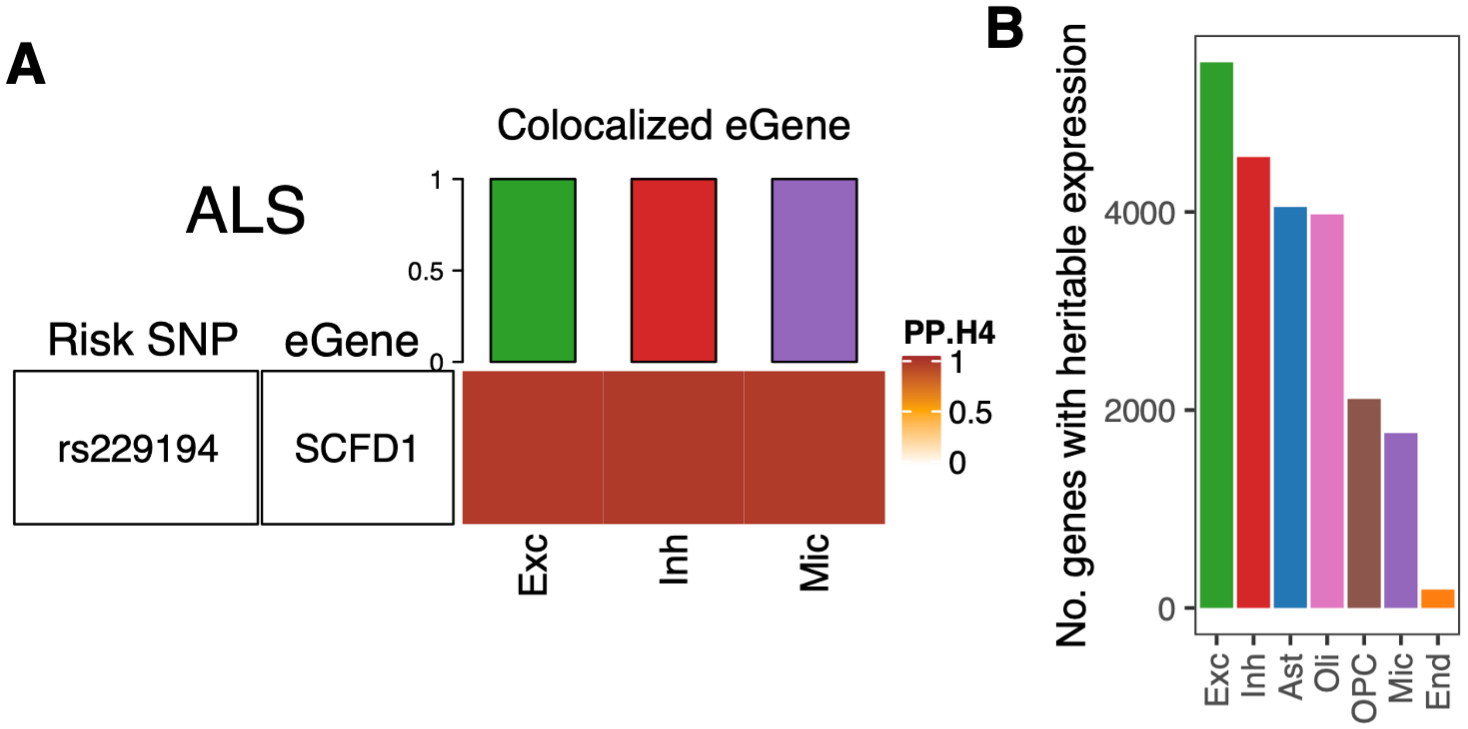

**Supplementary Figure 12. Association between eQTL and GWAS of neurodegenerative disease. (A) Colocalization of cell type-specific eQTL with risk SNPs of amyotrophic lateral sclerosis GWAS (van Rheenen *et al*., 2021). Elements of heatmap show posterior probabilities of the hypothesis H4 (PP.H4), which assumes GWAS and eQTL share a single causal SNP. Top bar chart shows the number of colocalized eGenes with PP.H4 > 0.8. Locus, locus name of the risk SNP. (B) Number of genes whose expression level was heritable. Expression heritability was assessed using the GREML software.**

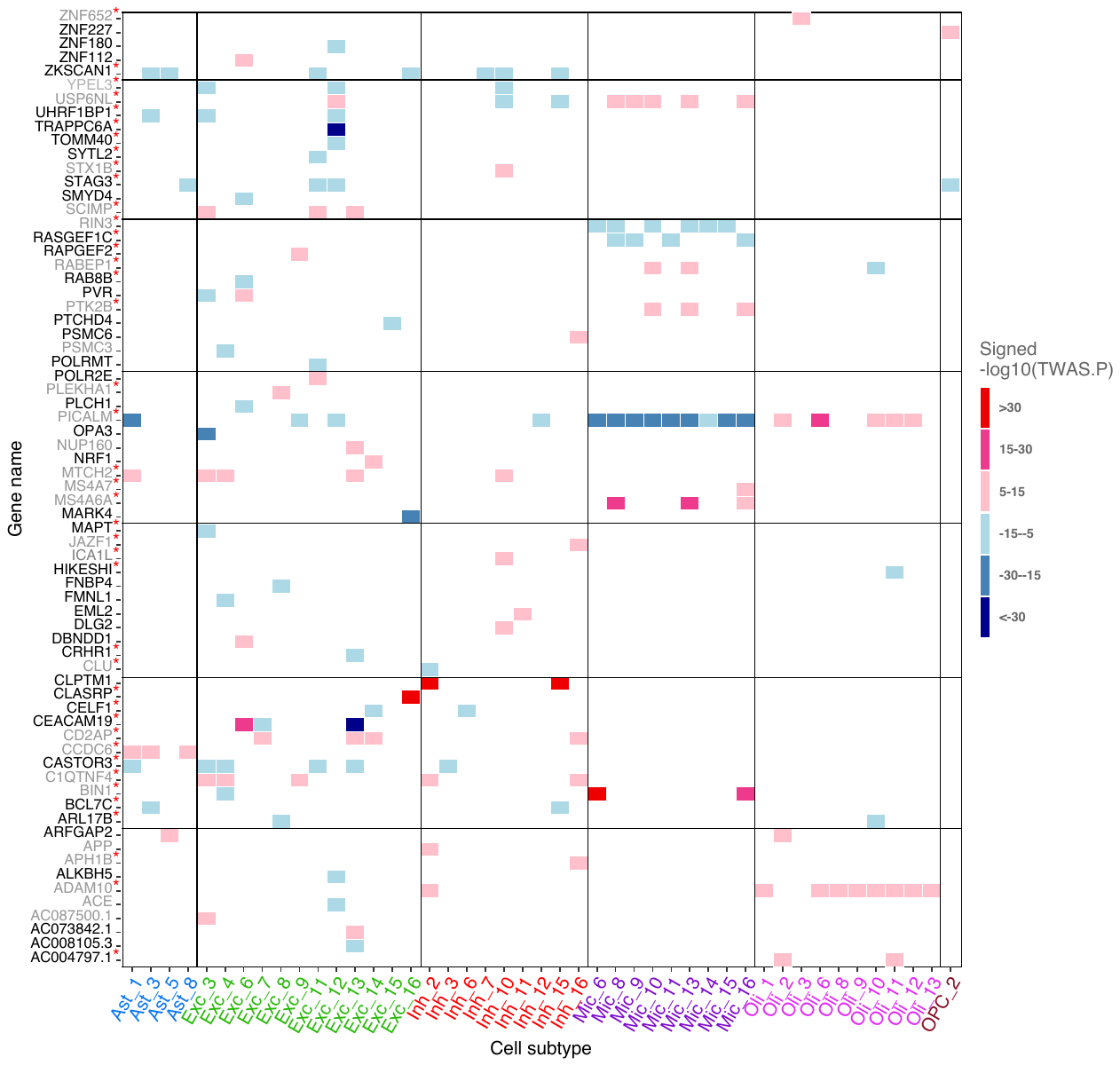

**Supplementary Figure 13. Cell subtype-level TWAS of Alzheimer's disease. Subtype-level pseudobulk expression and GWAS summary statistics of Wightman *et al*., 2021 were used as input. Non-gray elements show significant TWAS genes (FDR ≤ 0.05), and their colors represent effect directions and *p*-values. In row names, novel and known candidates for AD risk genes are colored black and gray, respectively. Red asterisk indicates that the same gene was also detected by cell type-level TWAS.**

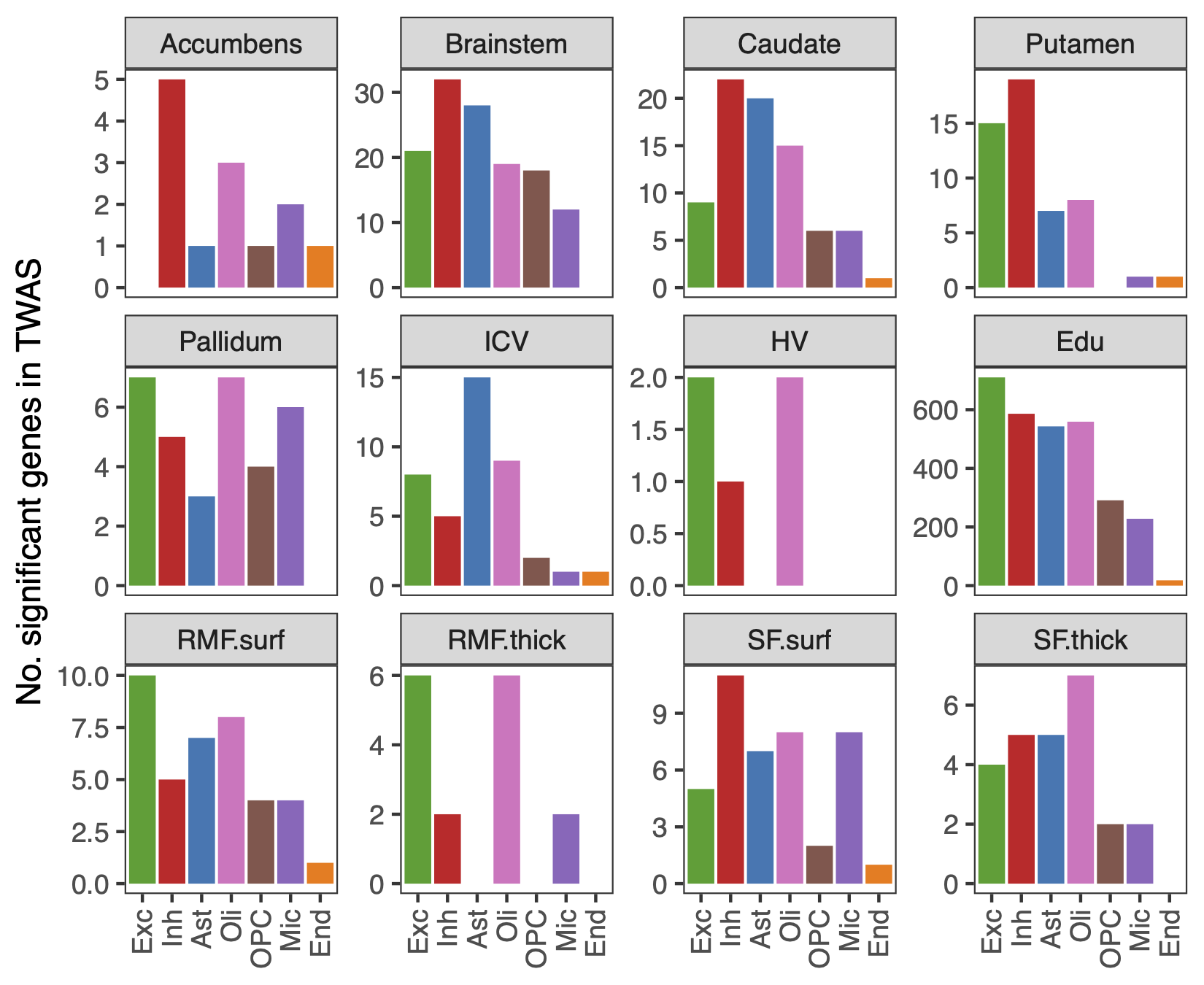

**Supplementary Figure 14. TWAS for brain volumes and educational attainment. ICV, intracranial volume; HV, hippocampal volume; Edu, educational attainment; RMF, the rostral middle frontal gyrus; SF, the superior frontal gyrus; RMF.surf, surface area of RMF; RMF.thick, cortical thickness of RMF; SF.surf, surface area of SF; SF.thick, cortical thickness of SF. Cell types are in order of descending expression heritability.**
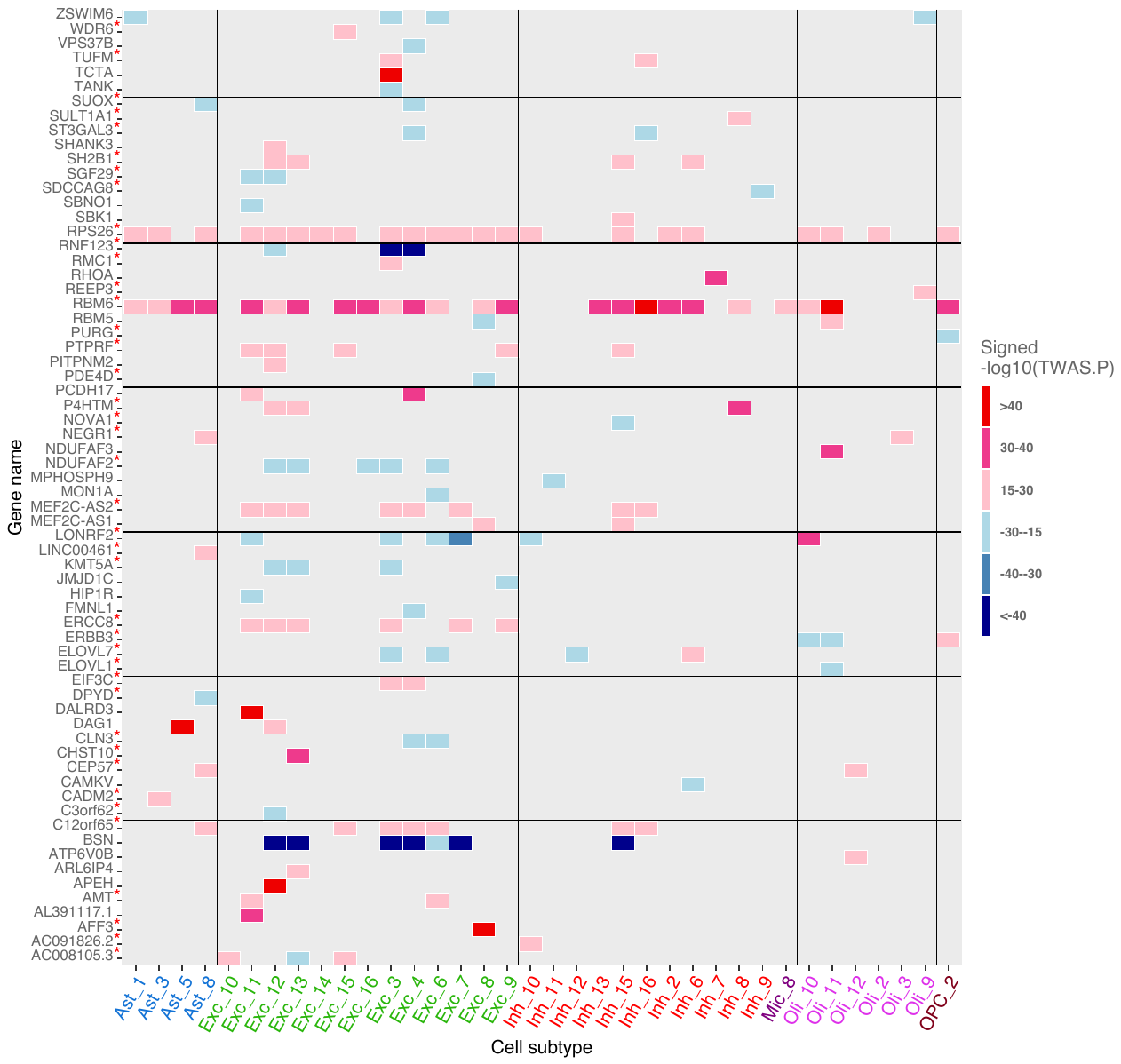

**Supplementary Figure 15. Cell subtype TWAS of educational attainment.** **Subtype-level pseudobulk expression and GWAS summary statistics of Lee *et al*., 2018 were used as input. Non-gray elements show significant TWAS genes (FDR ≤ 0.05), and their colors represent effect directions and *p*-values. In row names, red asterisk indicates that the same gene was also detected by cell type-level TWAS.**

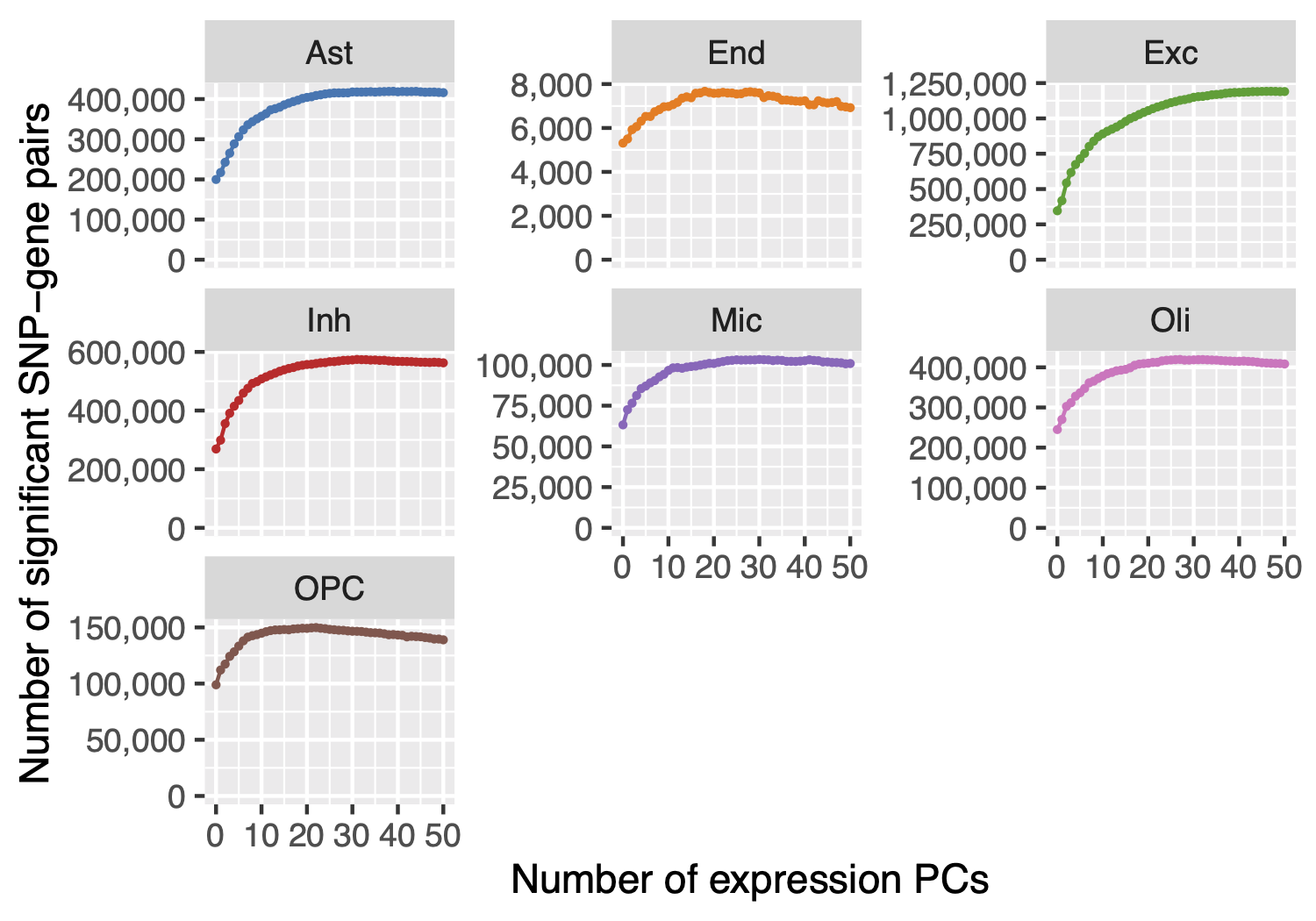

**Supplementary Figure 16. Number of eQTL as a function of number of expression principle components (PCs) that were used as covariates for linear modeling.**

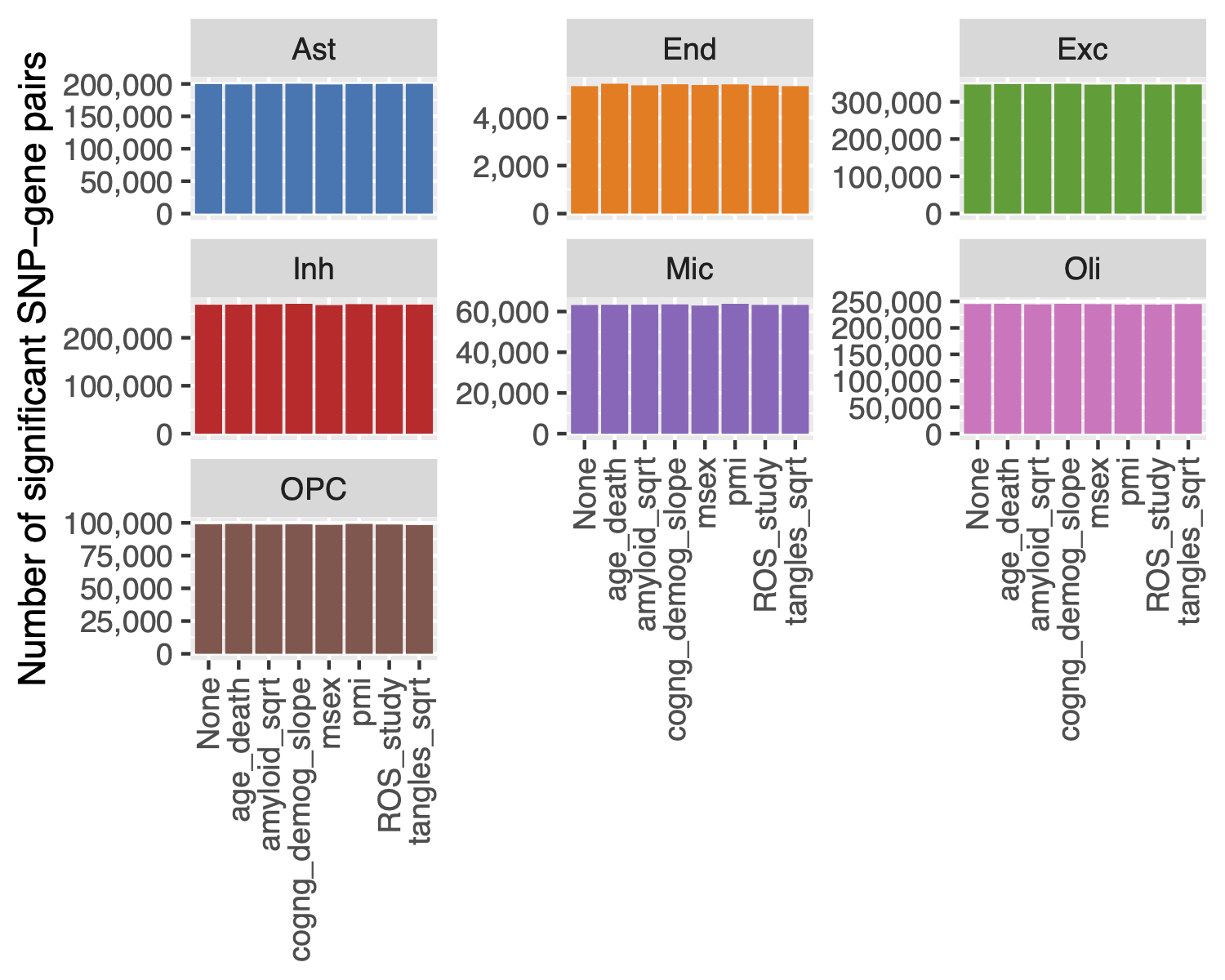

**Supplementary Figure 17. Dependency of number of eQTL on covariates of clinical traits. age_death, age at death; amyloid_sqrt, square root of overall amyloid level; cogng_demog_slope, estimated slope for global cognition controlled for demographics; msex, male sex; pmi, post-mortem interval; ROS_study, ROS or MAP cohort; tangles_sqrt, square root density of neuronal neurofibrillary tangles.**
